# Deciphering the spatio-temporal transcriptional and chromatin accessibility of human retinal organoid development at the single cell level

**DOI:** 10.1101/2023.07.19.549507

**Authors:** Birthe Dorgau, Joseph Collin, Agata Rozanska, Veronika Boczonadi, Marina Moya-Molina, Rafiqul Hussain, Jonathan Coxhead, Tamil Dhanaseelan, Lyle Armstrong, Rachel Queen, Majlinda Lako

## Abstract

Molecular information on the early stages of human retinal development remains scarce due to limitations in obtaining early human eye samples. Pluripotent stem cell-derived retinal organoids provide an unprecedented opportunity for studying early retinogenesis. Using a combination of single cell RNA-Seq and spatial transcriptomics we present for the first-time a single cell spatio-temporal transcriptome of retinal organoid development. Our data demonstrate that retinal organoids recapitulate key events of retinogenesis including optic vesicle/cup formation, formation of a putative ciliary margin zone, emergence of retinal progenitor cells and their orderly differentiation to retinal neurons. Combining the scRNA-with scATAC-Seq data, we were able to reveal cell-type specific transcription factor binding motifs on accessible chromatin at each stage of organoid development and to show that chromatin accessibility is highly correlated to the developing human retina, but with some differences in the temporal emergence and abundance of some of the retinal neurons.

## Introduction

Retinal organoids (ROs) generated from human pluripotent stem cells (PSCs) provide a powerful platform for studying retinal development, disease modelling, and cell-based replacement therapies [1]. Since their inception in early 2010s, multiple protocols have been developed by our group and others [2–5], resulting in laminated ROs with an apical putative outer nuclear layer packed full of photoreceptors, an inner nuclear layer comprising the retinal interneurons and a basal layer where retinal ganglion cells (RGCs) reside. Improvements in culture methods have led to the formation of a dense brush border comprising photoreceptor outer-like segments [6], which respond to light in a manner similar to postnatal mice at the eye-opening stage [7–9]. Single cell (sc) RNA-Seq and immunofluorescence analyses have shown the presence of all the retinal cell types within the organoids, which transcriptionally convergence to adult peripheral retinal cell types [8, 10]. While the development rate of organoids *in vitro* and fetal retina *in vivo* is similar, the molecular and cellular composition of the organoids is highly dependent on the differentiation protocol [11].

The dynamics of chromatin accessibility plays a critical role in regulating cell fate specification and maturation during development. The assay for transposase-accessible chromatin with high throughput sequencing (ATAC-Seq) has been employed to reveal the chromatin accessibility of pluripotent stem cell derived ROs. Notably these studies have shown that chromatin accessibility dynamics recapitulates human retinogenesis, but some divergent features were also identified [12]. For example, the accessible chromatin region near the transcription factor Ptf1a, which determines horizontal and amacrine cell fates during retinal development [13], is different between fetal retina and ROs, which could be in part responsible for the reduction in amacrine cells in the latter [14]. Equally, an *RHO* enhancer, which is present in mature human rods, is missing in the RO rods [15]: this could be the determining factor for the more modest expression of RHO in the organoids compared to human retina.

Spatial localisation and cellular interactions are key determinants of cell fate and behaviour. In ROs retinal lamination and positioning of cell types in a manner that allows functional connectivity is essential to recapitulate the physiological functions of adult retina. Current single cell studies to date lack single cell spatiotemporal resolution and thus it has not been possible to visualise the very rapid changes that characterise transitions from optic vesicle stage to optic cup or the emergence of the retinal neuroepithelium. While immunofluorescence studies indicate retinal progenitor cell (RPCs) localisation within the apical layer of ∼1 month old organoids [16], we are unsure where RPCs and neurogenic progenitors derived therefrom are first localised and whether a putative ciliary margin (pCM) is present in ROs. To fill this gap, we have employed for the first-time spatial transcriptomics (ST) in combination with scRNA– and ATAC-Seq analyses in a time series of ROs comprising 8 time points from day 10 to day 210 of differentiation. This integrated analysis has enabled us to reconstruct the single cell spatio-temporal transcriptional and chromatin accessibility of ROs development, revealing the precise order of retinal neurons differentiation and their localisation, which closely mimics human retinogenesis, but with some differences in the temporal emergence and abundance of some of the retinal cell types. Our data provide compelling evidence for the presence of pCM at the very edge of ROs, with potential to generate early RPCs. Our work provides the first integrated molecular and spatial atlas of human ROs development that could be used to identify novel genes and key pathways that underpin human retinal development and function.

## Results

### Single cell spatiotemporal phenotyping of ROs development

Eye-field specification, optic cup formation and RPCs emergence present important milestones for human retinal development, however these events are often difficult to study due to scarcity of human embryonic material. ROs derived from human pluripotent stem cells (PSCs) have been shown to recapitulate the early retinal development [17] and were used herein to generate a single cell spatio-temporal atlas up to day 210 of development using a combination of single cell (sc) RNA-and ATAC-Seq, and ST approaches. ROs were categorised according to age in following developmental stages: early retinal development (day 10, 20 and 35), mid retinal development (day 45, 60 and 90) and late retinal development (day 150 and 210).

### Early retinal development

The early retinal development stage in ROs was commonly characterised by two key events: the eye field/optic vesicle formation and the emergence of RPCs/neurogenic RPCs (NRPCs) occurring in a sequential manner (**Fig. 1**). The first time point, day 10 of differentiation, was dominated by optic vesicle epithelial clusters, revealing 73% and 87% abundance in scRNA and ST samples, respectively (**Fig. 1A, D, G** and **Fig. S1A**). These clusters were characterised by high expression of several classical eye field transcription factors such as *PAX6, NR2F1 and OTX2* (**Table S1**, **S2**; [18, 19], and typical optic vesicle markers including *FZD3, IGFBP5* and *VIM* [19–21], and covered nearly the whole of organoids (**Fig. 1G, D** and **Fig. S1A**). Cell proliferation was a common feature of several optic vesicle clusters as demonstrated by high expression of *MKi67, TOP2A, TYMS, HELLS, MCM4* and *HIST1H4C* (**Table S1**).

**Figure 1:**
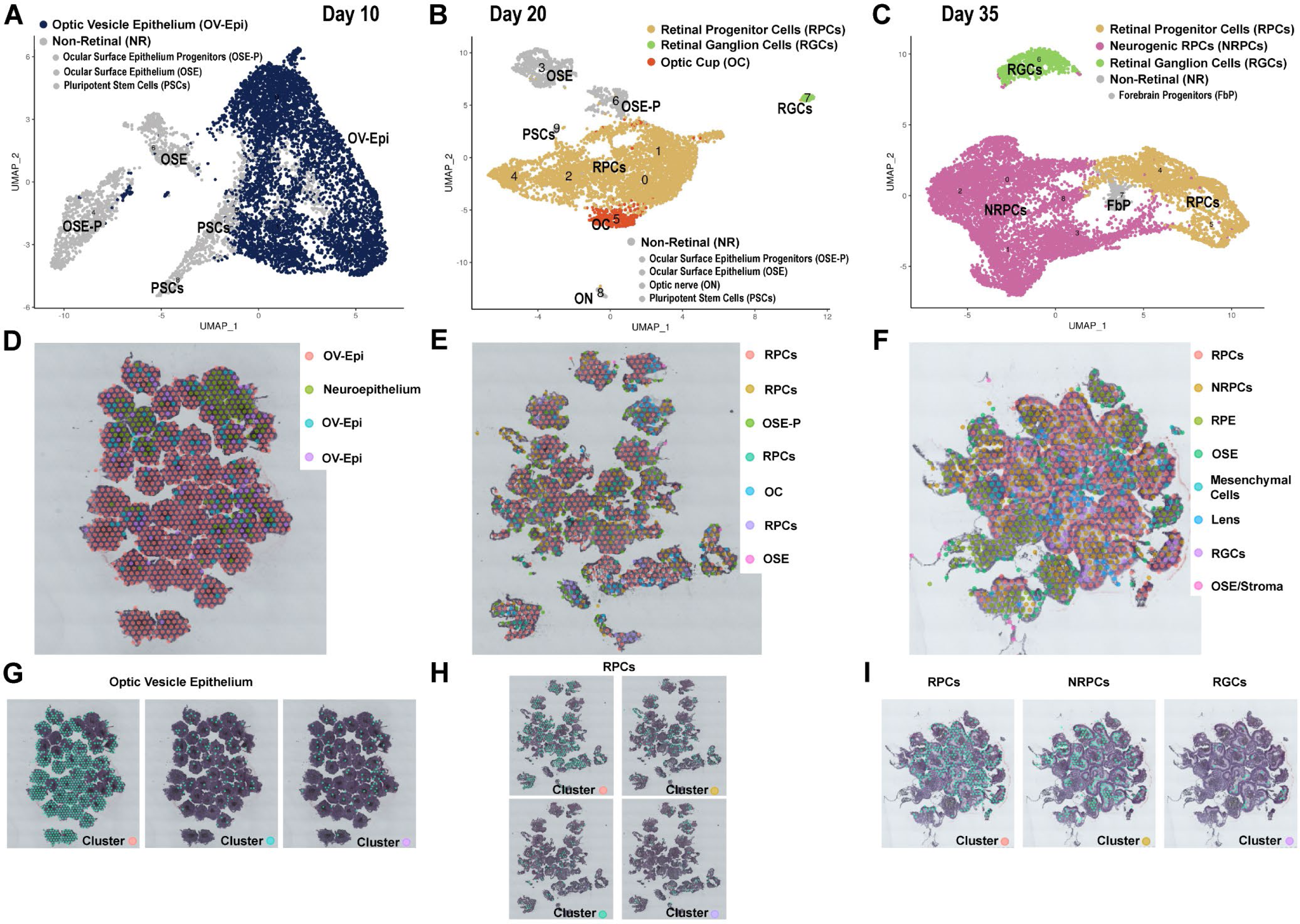
Early retinal organoid development (day 10 – 35) reveals key features of human retinogenesis, starting with optic vesicle formation and followed by the emergence of RPCs and RGCs. **A-C**) UMAP plots of scRNA-Seq in ROs at day 10 (**A**), day 20 (**B**) and day 35 of differentiation (**C**). **D-I**) Spatial localisation of the individual clusters in ROs at day 10 (**D, G**), day 20 (**E, H**) and day 35 (**F, I**) of differentiation acquired from ST analyses. Each cluster was identified based on expression of retinal specific cell markers shown in Table S1 for scRNA-Seq data and in Table S2 for ST data.

As expected, a small cluster of undifferentiated PSCs was still present in day 10 ROs (**Fig. 1A** and **Table S1**). Notably, two clusters of ocular surface epithelium (OSE), characterised by high expression of *KRT8, KRT18* and *KRT19*, were identified by scRNA-Seq, indicating that the culture conditions used for generation of ROs were also permissive to emergence other important eye related cell populations such as OSE cells (**Fig. 1A** and **Table S1**).

Optic vesicle epithelium developed over time, and by day 20 of RO differentiation, only a small cluster (7.6% of total cells) of optic cup cells characterised by high expression of eye field and pigmentation markers (*MITF, VAX2, PMEL, LHX2, PAX6)* was found alongside a large dominant heterogenous pool of RPCs (**Fig. 1B, E, H, Fig. S1B** and **Tables S1, S2**). RPCs were characterised by high expression of well-established markers including *ID3, ID1, SFRP2, VSX2, PAX6 and SIX3* [17] and represented 74% of total cells (**Fig. S2A** and **Table S1**, **S2**). At this stage of differentiation, RPC clusters revealed a broad distribution and were located all over retinal organoids without any specific layering and/or location (**Fig. 1E, H**). A small but distinct retinal ganglion cell (RGC) cluster characterised by highly expressed genes such as *GAP43, SNCG, PRHP, STMN, ELAVL3, ELAVL4, NEFL, NEFM, TBR1* was detected as early as day 20 of differentiation (**Figure 1B** and **Table S1**), which is slightly earlier than previously shown [17, 22]. This RGC cluster was only apparent in the scRNA-Seq data set due to either the current resolution of ST dots (50 µm diameter with a 100 µm centre to centre distance between spots) or the scarcity of this cluster. The RGC population increased over time, enabling the detection by both methods at day 35 of differentiation (**Fig. 1C, F, I** and **Fig. S1C, S2B**) and revealing their location at the basal side of ROs (**Fig. 1F, I**). The most striking feature of this differentiation time point was the increased pool of neurogenic retinal progenitor cells (NRPCs), especially noted in scRNA-Seq analysis (**Fig. 1C** and **Fig. S2B**). NRPCs (cluster 1) were characterised by high expression of *NEUROD2, NEUROD6, SOX4, SOX11, HES6, GADD45G* and displayed a distinct spatial profile to RPCs as revealed by ST analysis (**Fig. 1F, I, Fig. S1C** and **Table S2**). RPCs were found in an organised layer at the organoid’s apical edge, while NRPCs were predominantly located in the basal side (**Fig. 1F, I**). Other non-retinal cell clusters including OSE progenitors, OSE, OSE/Stroma, mesenchymal cells, lens, and forebrain cells were identified in day 20 and/or 35 (**Fig. 1, Fig. S1B-C** and **Tables S1**, **S2**).

In summary, single cell RNA-Seq and ST demonstrate that the early development in ROs recapitulates the key events known from morphological and immunofluorescence studies of retinal development, starting with the eye field specification/optic vesicle formation, the emergence of RPCs and NRPCs, followed by the birth of an early born retinal cell type, the RGCs.

### Mid retinal development

The mid developmental stage of retinal organoids was characterised by the reduction of RPCs and NRPCs pool over time and changes in their spatial localisation (**Fig. 2** and **S2A**). Whilst RPCs were mainly located at the apical side of retinal organoids at day 45 and 60 (**Fig. 2D, E, G, H**), NRPCs were found more basally towards the centre of organoids at day 90 of differentiation (**Fig. 2F, I**). Notably, we identified specific clusters (cluster 7 in the scRNA-Seq and cluster 4 in the ST analyses at day 45, and cluster 8 scRNA-Seq of day 60) (**Fig. 2A, B, D and Fig. S1D**) that were characterised by high expression of ciliary margin (*WNT2B*), eye field (*PAX6),* pigmented cell (*PMEL, TYRP1)* and RPCs (*ID3, ID1, SFRP2, ZIC1)* markers (**Tables S1, S2)** [17, 23]. In a recent spatiotemporal phenotyping single cell analysis, we have observed the very same transcriptional signature in the ciliary margin zone of developing human retina and have shown that this region transiently harbours the early RPCs [24]. Using immunofluorescence and pulse labelling studies, Kuwahara and colleagues demonstrated the presence of a ciliary margin like stem cell niche in ROs, comprising a continuous neural retina epithelium with a small, pigmented domain of adjacent retinal pigment epithelium (RPE), which displayed the expression of both neural retina and RPE markers, with the capacity to expand the neural retina by *de novo* progenitor generation [2]. Based on these findings we defined the clusters with high expression of ciliary, pigmented and RPCs markers, as putative ciliary margin (pCM). In accordance, ST analysis revealed the location of this cluster at the very edge of day 45 ROs, mimicking the position of the ciliary margin zone in the developing human retina (**Fig. 2G**).

**Figure 2:**
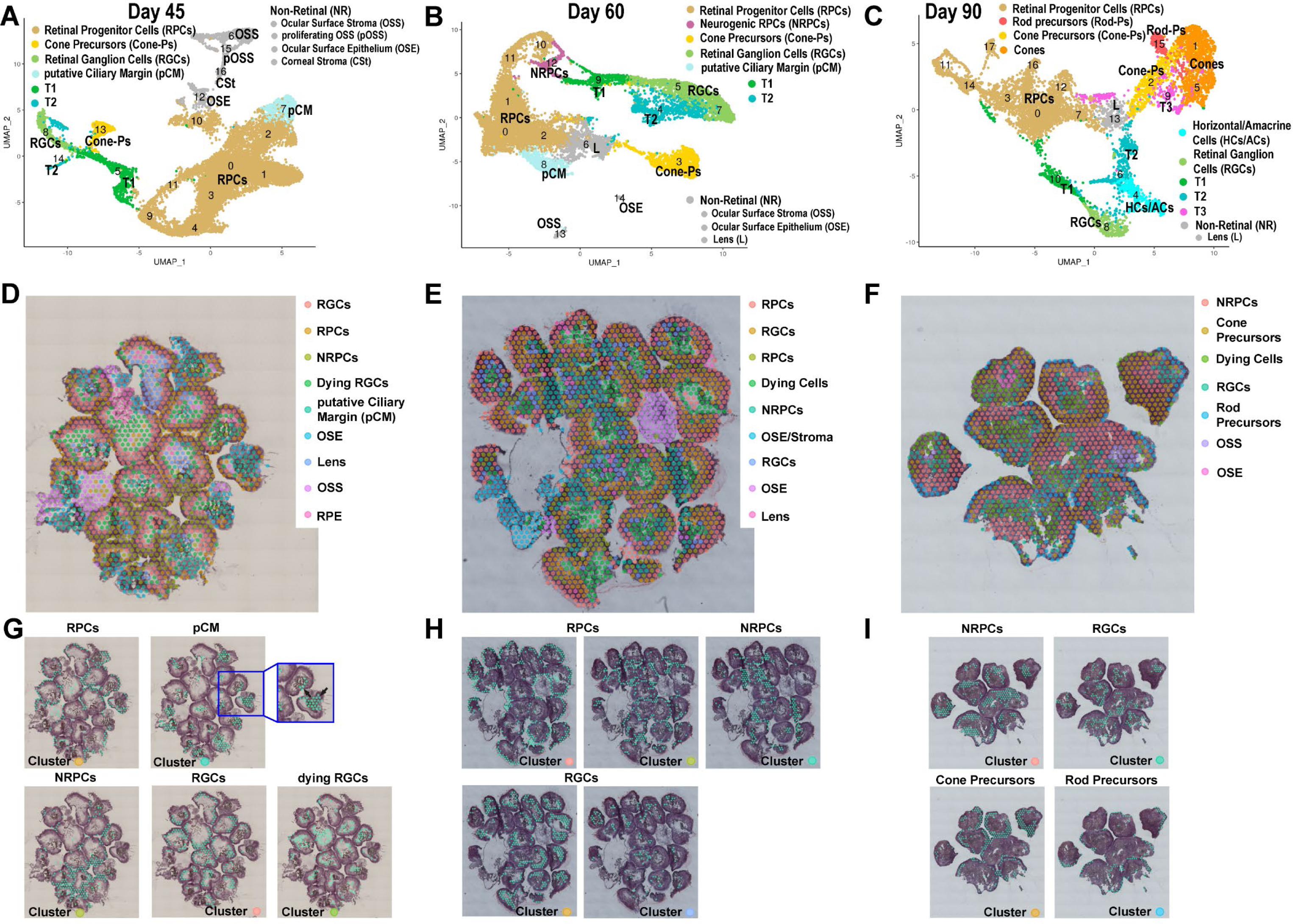
Mid organoid development (day 45-90) reveals the presence of pCM and specific localisation of RPCs, NRPCS, RGCs and photoreceptor precursors. **A-C**) scRNA-Seq UMAPs of hiPSC-derived ROs at day 45 (**A**), day 60 (**B**) and day 90 (**C**). **D-I)** Individual cluster location in ROs at day 45 (**D, G**), day 60 (**E, H**) and day 90 (**F, I**) of differentiation identified from the ST analyses. Cluster annotation is based on highly expressed retinal cell key marker gene expression shown in Table S1 for scRNA-Seq data and in Table S2 for ST data.

The ST analyses of mid retinal development did not distinguish between the various types of NRPCs, however, the scRNA-Seq revealed the emergence of three transient T1, T2 and T3 NRPCs, which have been shown to give rise to defined types of retinal neurons at specific developmental windows during human retinogenesis (**Fig. 2A-C** and **Table S1**; Sridhar, Hoshino et al. 2020). T1 cluster was located between the RPCs and retinal neurones and was defined as early as day 45 of RO differentiation (**Fig. 2A** and **Table S1**). A T2 cluster was found next to T1 and RGCs at day 60 of differentiation, but at day 90 the very same cluster was found adjacent to a mixed population of horizontal/amacrine cells (**Fig. 2B, C**). The T3 cluster became apparent at day 90 of RO differentiation and was located at the base of emanating cone and rod photoreceptor precursors clusters (**Fig. 2C** and **Table S1**).

The RGCs clusters, the second largest besides RPCs/NRPCs in the mid-developmental stage reached its peak at day 60 and then decreased from this day onwards, reflecting the increasing cell type complexity of ROs (**Fig. 2B, E, H** and **Fig. S2B**). Strikingly, our ST analyses provided evidence of dying RGCs at day 45 of RO differentiation (**Fig. 2D** and **Fig. S1D**): these were characterised by the expression of many mitochondrial and ribosomal related genes in addition to RGCs marker genes such as *SNCG, NEFL, DCX, STMN2* (**Table S2**). Those dying RGCs were located in the centre of retinal organoids (**Fig. 2D, G**), reflecting the drastic reduction of RGCs during human retinal development. Dying cells located in the middle or organoids were also observed at day 60 and 90 of RO differentiation (**Fig. 2E, F**), corroborating the presence of an apoptotic core often visible by bright field microscopy in ROs.

The first photoreceptor precursors to emerge were cone precursors: these were observed as early as day 45 of differentiation (cluster 13; **Fig. 2A**) and expressed at high level cone-specific genes such as *THRB, CRX, PDE6H, PRDM1, DCT, IMPG2, OTX2* as well as some RGC-specific (*SNCG, DCX, STMN2*), horizontal cell specific (*ONECUT1, ONECUT2 and ONECUT3*), and NRPCs-specific (*ATHO7, NEUROD1, VXN, GADD45G, SCG, PCP4*) markers (**Table S1**). Similar cone cell clusters co-expressing RGC and interneuron markers were recently identified by our group in the retinas of 8-13 PCW of human specimens [24]. This highlights the dynamic transcriptional profile of photoreceptor precursors during retinogenesis and points to the immature state of cones at this time of differentiation. The expression of NRPCs, RGCs and interneuron-related genes dissipated from cone precursor’s transcriptional profile at day 60 and 90 of RO differentiation, and cone precursor clusters were characterised by high expression of cone typical markers such as *PDE6H, ARR3* and *THRB* (**Table S1**, **S2**). Cone precursors were observed spatially within retinal organoids from day 90 of differentiation onwards, forming a thick organised layer at the apical side of ROs (**Fig. 2F, I**). Rod precursors first identified at day 90 by both methods formed a thin layer above the cone cluster (**Fig. 2C, F, I**) and were characterised by highly expressed genes such as *NRL, RHO, ROM1, NR2E3, PDE6G, PDE6A, GNGT1, GNAT1, GNB1* (**Tables S1**, **S2**).

Apart from ROs containing cell clusters described above, a different type of organoid was apparent from approximately day 30 of differentiation onwards, revealing a small retinal-like core with a big opaque cystic-like structure attached (**Fig. S3A**), shown to contain ocular surface stroma (OSS) and OSE cell clusters by the ST analyses (**Fig. 2D-E, Fig. S1D-E** and **Table S2**). These organoids were named ‘eye-like organoids’ hereafter and were further investigated using scRNA-Seq and/orimmunofluorescence (IF) analysis at day 45 and day 90 of differentiation respectively (**Fig. S3** and **Table S3**). Strikingly, all corneal cell types forming the three layers of the cornea were identified: the endothelium, the stroma and the epithelium, which was further defined as conjunctival and corneal-limbal epithelium at day 45 (**Fig. S3B**), and then further stratified to superficial corneal-limbal epithelium and basal corneal/conjunctival epithelium at day 90 (**Fig. S3C**). The corneal endothelium cluster was characterised by high expression of *TWIST1, TWIST2, BMP4, PAX3, PDGFRA and TAGLN* genes, while genes such as *LUM, COL1A1, COL1A2, COL3A1, COL5A1, COL6A2, OGN, POSTN, FOXC1, FBLN1* were highly expressed in the corneal stoma cluster (**Table S3**). The conjunctival and corneal-limbal epithelial clusters were characterised by high expression of *KRT13* and *KRT15* respectively (**Table S3**). Retinal cell clusters were also present including pCM, RPCs, RGCs and cones as well as T1 at day 45 and a mixed cluster of HC/ACs at day 90 (**Fig. S3B** and **Table S3**).

IF analysis at the “eye-like organoids” at day 45 of differentiation confirmed the presence of conjunctival epithelium marked by the expression KRT13, limbal-corneal epithelium characterised by the expression of KRT15 at the apical edge of corneal organoids, and corneal stroma cells by the expression of lumican (LUM) inside the organoids (**Fig. S3D**). At this time point many proliferative epithelial progenitor cells immunostained with MT2A and Ki67, and Np63, were detected within the eye-like organoids (**Fig. S3D**). Alongside the corneal clusters, the organoids contained a small retinal core, showing thick Crx-positive rosette-like structures (**Fig. S3D** and **Table S3**). The scRNA-Seq also revealed the presence of lens clusters at day 90 in the “eye-like organoids” (**Fig. S3C**), corroborated by the presence of CRYAA and CRYAB immunostained cells within the “eye-like organoids” (**Fig. S3D**). The “eye-like organoids” didn’t develop further and were very rarely found after day 90, most likely due to an inappropriate differentiation media which was tailored for the neuronal retinal differentiation.

In summary, the mid-developmental stage was the most dynamical stage in where RPCs/NRPCs develop to transient cell progenitors, which then gave rise to retinal cell types such as interneurons and photoreceptor precursors. Notably, another type of organoid containing corneal, lens and retinal cell specific clusters were present at this stage, but their further survival was compromised by the culture media optimised for ROs.

### Late retinal development

The late developmental stages were characterised by the smallest pool of RPCs/NRPCs in comparison to other stages and an increasing population of mid to late born retinal cell types such as photoreceptors, bipolar (BCs) and Muller glia (MC) cells (**Fig. 3, Fig. S1G, H** and **Fig. S2C, D**). ST demonstrated the presence of remaining RPCs within the pCM clusters (**Fig. 3E, F** and **Table S1, S2**), which were characterised by high expression of ciliary muscle (*CPAMD8, COL9A2, ACTC1, MYL1*) [25] in addition to RPC markers (*ZIC2, SFRP2, IDE1, ID3*) at day 150 and 210 of RO differentiation (**Table S2**). The T2 transient neurogenic cluster, which gives rise to horizontal and amacrine cells, was not present at this late stage of differentiation in accordance with the earlier emergence of these interneurons during the mid-stage of organoid development (**Fig. 3A, B**). The T1 neurogenic progenitors which give rise to T3 and further differentiate into bipolar cell precursors (BCPs) and photoreceptors were identified by scRNA-Seq at day 150 of differentiation but not at day 210 (**Fig. 3A, B**), in accordance with the increased presence of photoreceptors and detection of BCPs during the late development stage (**Fig. 3 and S2C, D**).

**Figure 3:**
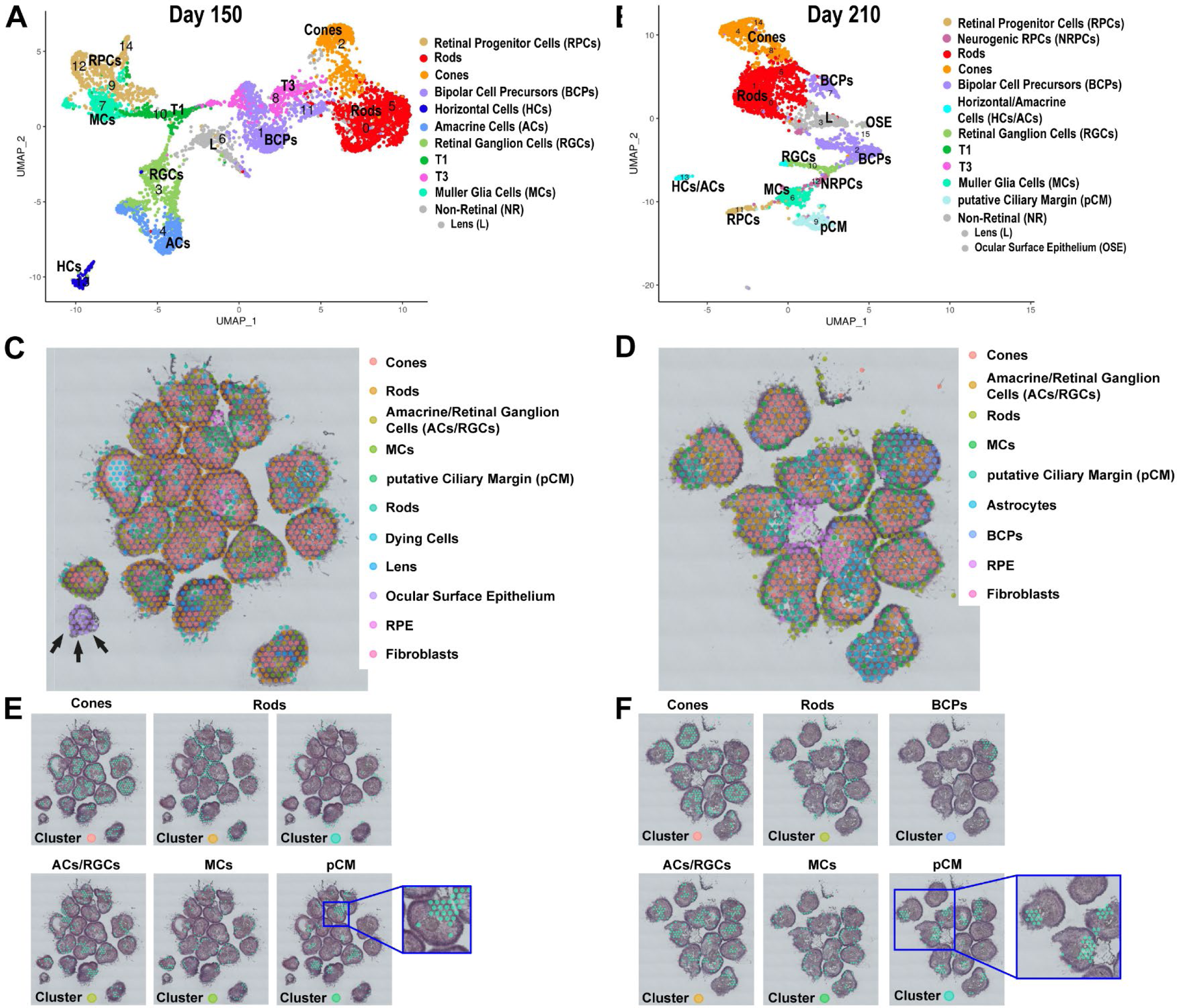
Late development of ROs (day 150 – 210) reveals maturation of photoreceptors, the appearance of BCs and MCs. **A, B**) UMAP plots of scRNA-Seq of ROs at day 150 (**A**) and day 210 of differentiation (**B**). **C-F)** Localisation of individual clusters in ROs at day 150 (**C, E**), day 210 (**D, F**) of differentiation based on ST analyses. Each cluster was identified based on expression of retinal specific cell markers shown in Table S1 for scRNA-Seq data and in Table S2 for ST data.

BCPs and Muller Glia cells (MCs) were first identified at day 150 of differentiation and were the last retinal cell type to emerge in retinal organoids (**Fig. 3**). This is consistent with birth dating and bulk RNA-Seq studies, stating BCs and MCs to be of the latest cell types developing during retinogenesis [26, 27]. MCs were detected by both methods and were characterised by high expression of *RLBP1, GPX3, CRYM, CLU, DIO3, SLC1A3* (**Table S1**, **S2**). They were found in between the photoreceptor layer at the apical edge of organoids (**Fig. 3E, F**), reflecting their apical processes, which form tight junctions with each other and/or inner segments of photoreceptors, establishing the outer limiting membrane. scRNA-Seq data indicated the close relationship between MCs and RPCs clusters (**Fig. 3A, B**) corroborating the MCs origin from RPCs shown earlier by Sridhar and colleagues [22]. BCPs comprised 15% of total cells at day 150 and day 210 of differentiation (**Fig. S2D**) and were detected by ST in the outer layer of ROs (**Fig. 3D, F**).

Rods and cone photoreceptor clusters, first detected in the mid developmental stage, increased in occurrence (**Fig. S2C**) and matured by developing inner/outer segments during the late development stage (**Fig. S4G, H**). ST showed that rods were located close to the brush border of ROs and cones underneath those in the presumptive ONL (**Fig. 3C-F**). Overall, ROs from day 150 and day 210 of differentiation revealed a comparable cellular composition, although some populations such as rod photoreceptors increased in occurrence at day 210 (**Fig. S2**). Importantly, the scRNA-Seq demonstrated the presence of RGCs until the very last point of differentiation timepoint (day 210) (**Fig. 3B**).

Similarly, to mid stages of differentiation, other cell types including fibroblasts, lens, astrocytes and OSE cells were detected during the late differentiation (**Fig. 3** and **Fig. S1G, H**). ST demonstrated the presence of a few OSE organoids as a separate entity to ROs at day 150 of differentiation (black arrows in **Fig. 3C**), however these were not present any longer at day 210. Strikingly, astrocytes were detected in ROs of day 210 by ST (**Fig. 3D**). In summary, ROs in the late developmental stage contained all the retinal cell types and only a small pool of RPCs. Moreover, this stage was characterised by an expansion of late born retinal neurons such as rods, BCPs and MCs as well as the overall maturation of retinal cell types (e.g., photoreceptors developed inner/outer segments, **Fig. S4**).

### Pseudo-temporal analysis reveals a source of early RPCs in the putative ciliary margin and the transition of RPCs to neurogenic precursors and retinal neurons

Transcriptomes of retinal progenitors and retinal neurons from all developmental stages (day 10 – 210) were integrated to better understand the molecular events that guide specification of RPCs and their differentiation (**Fig. 4A**). After quality control, transcriptomes of 44,889 cells were obtained and merged, resulting in thirty-six transcriptionally distinct clusters visualised in UMAP. These clusters were defined using well-known and highly expressed marker genes for each retinal cell population (**Table S4**). Of those, sixteen clusters were identified as RPCs, two as NRPCs, four clusters as neurogenic transitional populations (T1, T2, T3), two clusters as pCM, one cluster as lens and eleven clusters as retinal neurons such rods, cones, bipolar cells, horizontal/amacrine cells, RGCs and MCs (**Fig. 4A** and **Table S4**).

**Figure 4:**
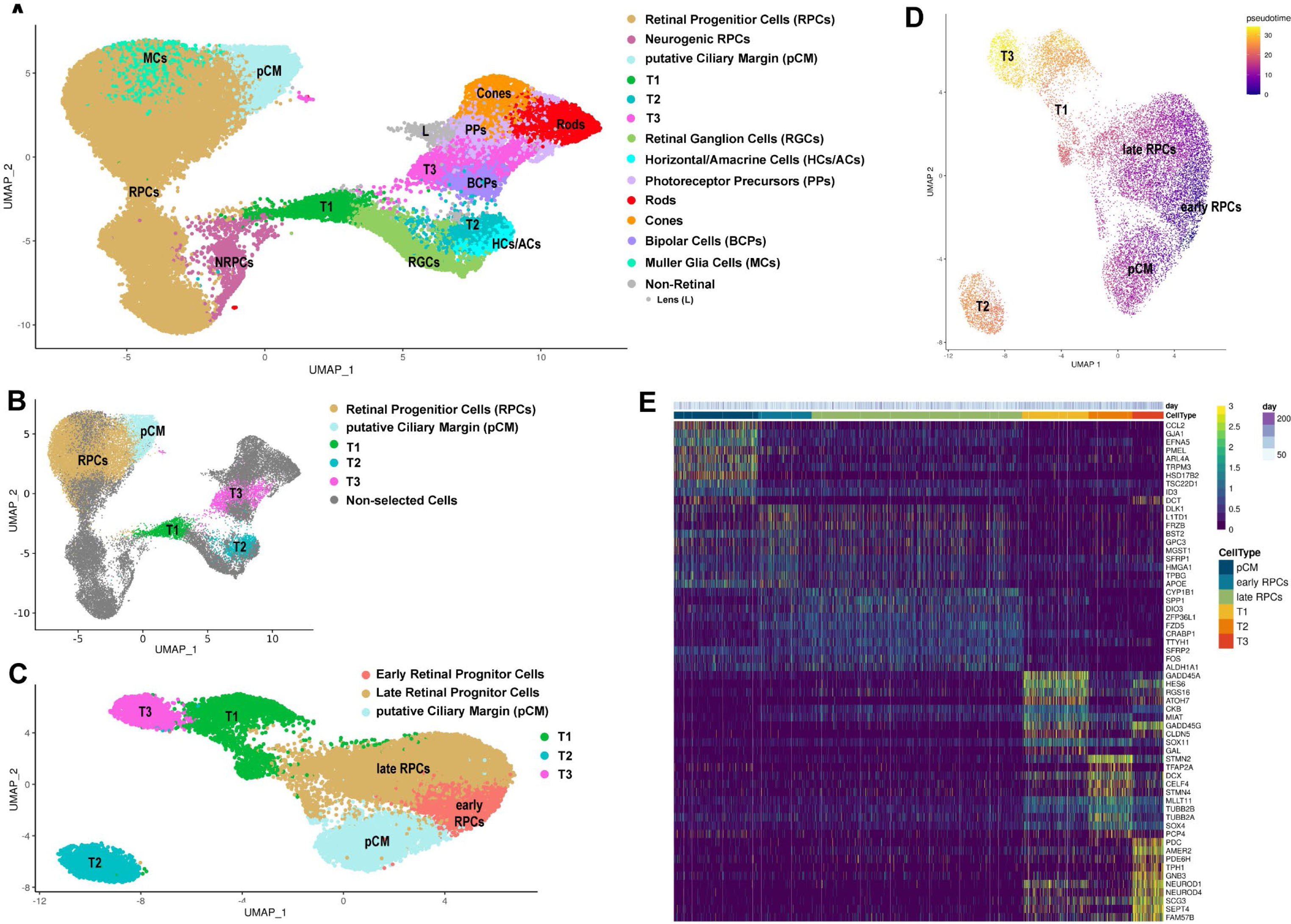
RPC identification and their developmental trajectories during RO development. **A**) UMAP plot of integrated scRNA-Seq of ROs during development from day 10 to day 210 of differentiation. Cluster annotations were based on highly expressed retinal key marker genes shown in Table S4. For ease of visualisation, clusters of the same cell type are shown by the same colour. **B)** Individual selected clusters of integrated scRNA-Seq retinal cells (**A**) used for pseudotime analyses. Highly expressed markers for each cluster along the pseudotime trajectory shown in Table S4. **C, D)** Pseudotime analysis demonstrating transition from pCM to early and late RPCs followed by T1 which further commits to either T2 or T3 transient neurogenic progenitors. **E)** Gene expression heatmap, indicating similarities in gene expression patterns between the pCM and early RPCs, and between T1 and T3 transient neurogenic progenitors.

To investigate the temporal transition of RPCs to neurogenic transitional progenitors T1, T2 and T3, pseudo-temporal analysis of gene expression changes was performed (**Fig. 4B-E**). Eight RPCs and two NRPC clusters characterised by high expression of genes such as *MKI67, TOP2A, UBE2C, CCND1, MCM3, UBE2C* and *CENPF* were excluded from the analysis because of their mitotic/proliferating nature (**Fig. 4B** and **Table S4**). As the pCM cluster possessed cells expressing early RPCs marker genes (**Table S4**), it was incorporated in the analysis (**Fig. 4B**). The RPCs and the RPC-expressing cells in the pCM cluster shared common marker genes, typical of RPCs such as *ID1, ID3, SFRP2, ALDH1A1* and *PCDH7*: in effect 28.3% of highly expressed genes in the pCM cluster were shared with early RPCs clusters. In addition, unique marker genes such as *ZIC1, FOXP1, PAX6* and *HSPB* were highly expressed in the pCM (**Fig. S5A** and **Table S4**). In particular, *ZIC1* (zinc finger protein of cerebellum 1) high expression was found mainly in the pCM cluster (**Fig. S5B**). RNA-Scope revealed the presence of *ZIC1* expressing cells exclusively at the very periphery of RO neuroepithelium (**Fig. S6A, B**), lying adjacent to the HES6 expressing cells that comprised the majority of retinal neuroepithelium, extending from the periphery towards the centre of retinal organoids (white arrowheads in **Fig. S6B**).

ZIC1 is a member of the ZIC family of C2H2-type zinc finger proteins which play a fundamental role in various early developmental processes. Mutations in members of the ZIC family have been associated with a wide variety of congenital malformations, including Dandy–Walker malformation, holoprosencephaly, neural tube defects, and heterotaxy [28, 29]. Koso and colleagues [30] showed that ZIC1 expression was restricted to SSEA1-positive RPCs in the peripheral region (ciliary margin) of the mouse retina, which has been postulated to contain stem cell like populations that can further develop to RGCs [31]. A similar stem cell niche with the potential to expand the neural retina was found in a ciliary margin like zone of retinal organoids [2]; hence, we hypothesise that the pCM cells that express RPCs markers may contribute to retinal neurogenesis, giving rise to early RPCs of the neural retina. This was corroborated by our pseudotime analyses, indicating the start of the developmental trajectory in pCM, leading to early RPCs (characterised by high expression of *ID3, MGST1, SFRP1, APOE1),* followed by late RPCs (characterised by high expression of *SPP1, DIO39, FOS, CRABP1* and *FZD5)* and the T1 transient neurogenic precursors (**Fig. 4C-E**, **Table S4**).

The T1 cluster led to two further transitional population (T2 and T3, **Fig. 4B**); in accordance, 13.6% of T1’s highly expressed markers were shared with T2 and T3 clusters (**Fig. S5C** and **Table S4**). RGCs characterised by high expression of *GAP43, PRPH* and *SNCG,* emanated from the transitional cluster T1 (**Fig. 5** and **Table S4**), confirming recent transcriptomic and birth dating studies of the human retina and ROs [22, 32, 33]. T2 gave rise to a mixed cluster of horizonal and amacrine cells characterised by high expression of *TFP2A, ONECUT1, ONECUT2, MEIS2*, *PAX6* and *GRIA4* (**Fig. 5** and **Table S4**).

**Figure 5:**
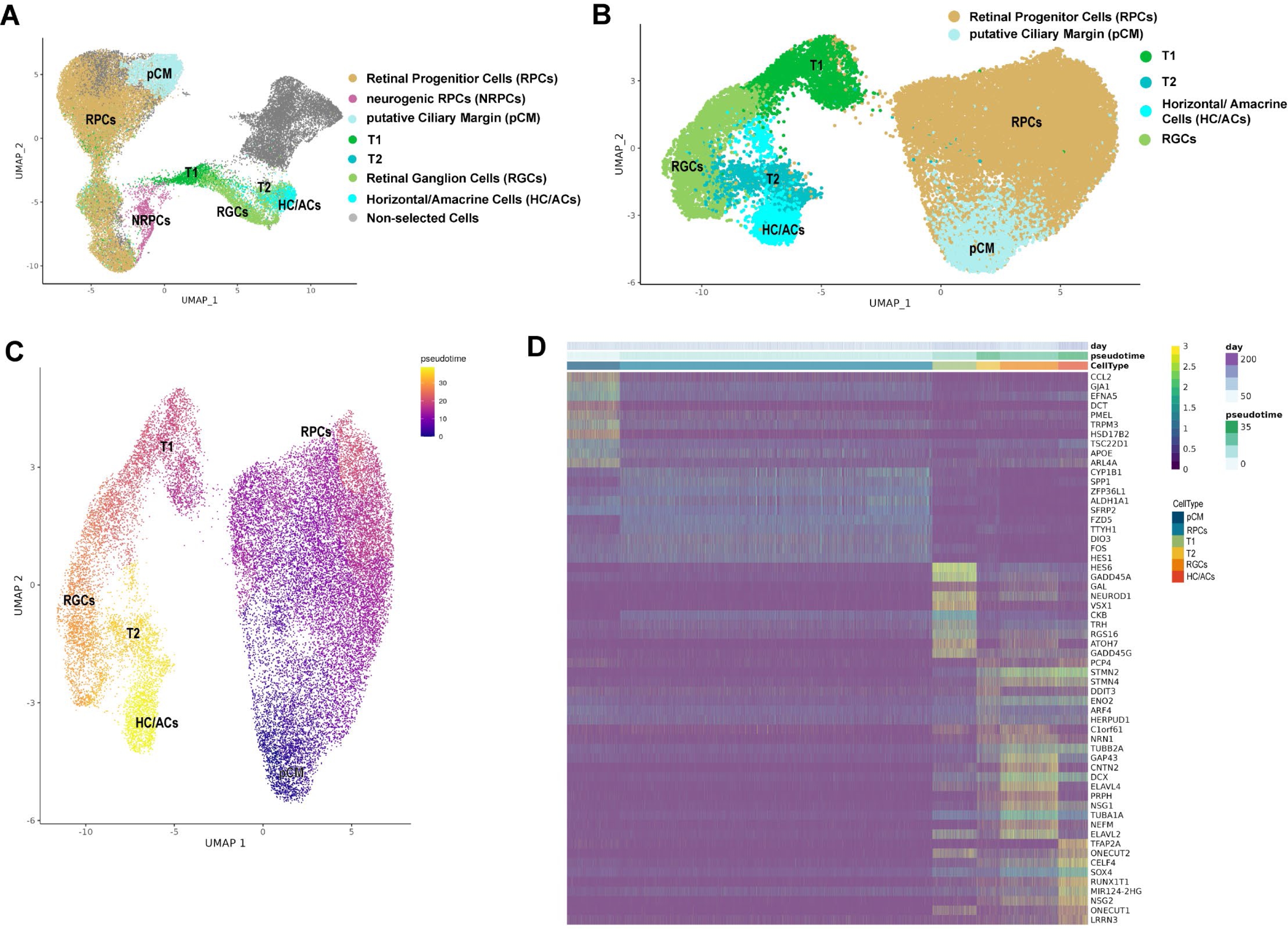
Pseudotime trajectory analysis indicates the emergence of horizontal and amacrine cells via T1 and T2 neurogenic progenitors. **A)** pCM, RPCs, T1 and T2 transient neurogenic cells and horizontal/amacrine cell clusters were selected from scRNA-Seq and used for pseudotime analyses. Key marker genes for each cluster are shown in Table S4. **B, C)** Pseudotime analysis revealing transition from pCM to RPCs, T1 and T2, further giving rise to horizontal/amacrine cells. **D)** Gene expression heatmap, showing some overlapping gene expression between T1 and T2 transient neurogenic progenitors as well as between T2 and RGCs, and horizontal/amacrine cell clusters.

Photoreceptor precursors and BCs originated from the transitional population T3 (**Fig. 6**), confirming previous studies (Sridhar, Hoshino et al. 2020). Although the BCs expressed several key marker genes such as *VSX1, NEUROD4, CADPS, PDRM8, VSX2* and *OTX2,* it also revealed strong transcriptional similarity to T3 (**Fig. S5D**). Based on this evidence, we concluded that ROs in this study contained BCPs, which explains their early temporal emergence (next to T3) in the pseudotime analyses (**Fig. 6B, C**). Cone precursors expressing *DCT, THRB, PDE6H, HES6, SCG3, NEUROD4,* NEUROD1, *VXN* and *CRX* genes led to mature cones, characterised by high expression of *PDE6H, ARR3, GUCA1A, GNGT2, GUK1, TULP1* and *AIPL1* (**Fig. 6D** and **Table S4**). Rod precursors expressing *NRL, RECOV, NR2E3, AIPL, NEUROD1, ROM1* and *CRX*, developed slightly later than cone precursors and matured over time to rods (**Fig. 6B-D**), which were characterised by high expression of genes such as *PDE6G, ROM1, GNGT1, GNAT1, NR2E3, RHO* and *CNGB1* (**Table S4**).

**Figure 6:**
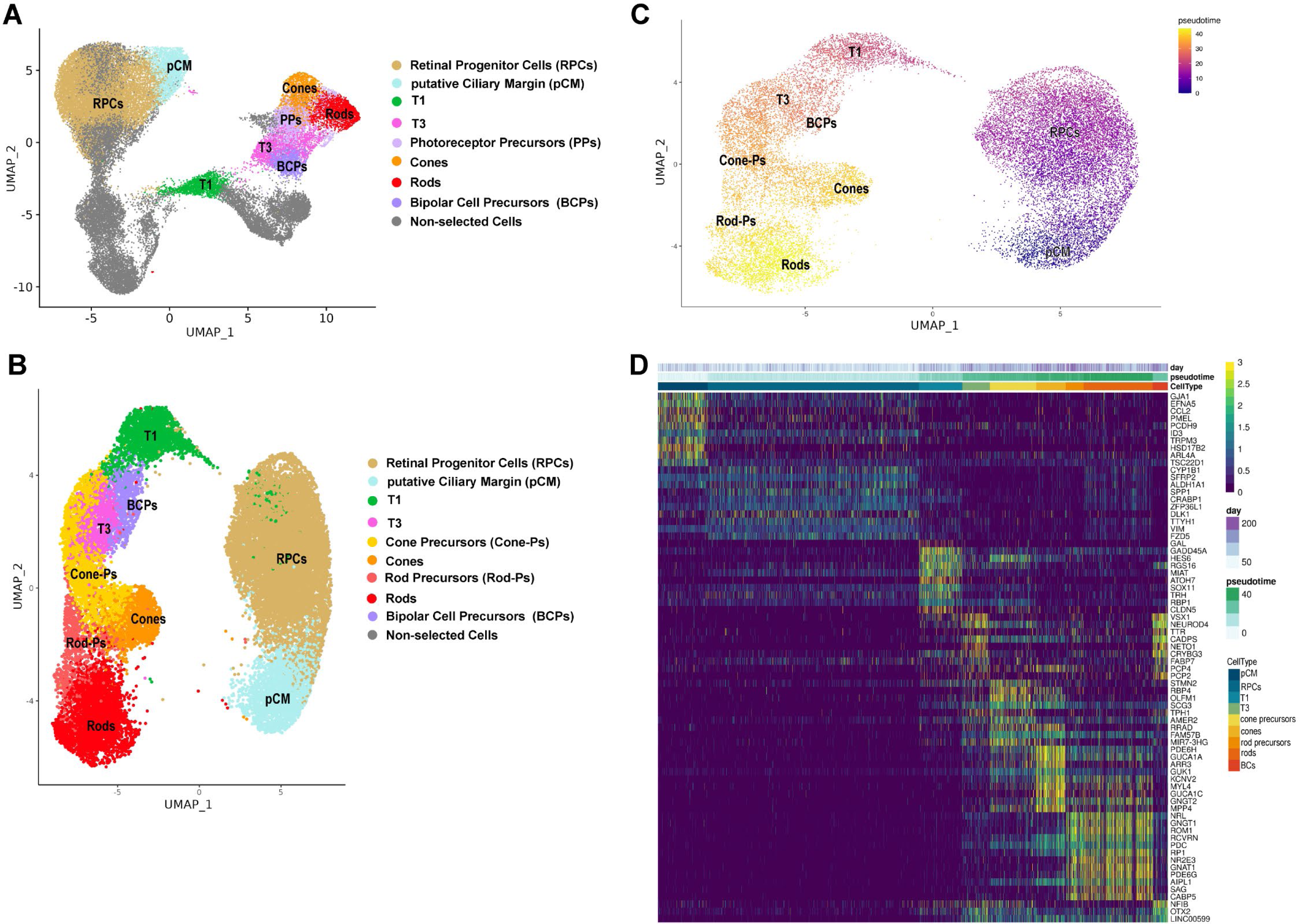
Emergence of photoreceptor and bipolar cells via T1 and T3 neurogenic progenitors. **A)** UMAP plot showing selected cluster of integrated scRNA-Seq (Table S4) used for pseudotime analyses. Highly expressed marker genes for each cluster along the pseudotime trajectory are shown in Table S4. **D)** Gene expression heatmap revealing distinct expression patterns of T1 and T3 progenitors and photoreceptors, while BCPs display transcriptional similarity with transient neurogenic cells, T1 and T3, suggesting the lack of mature bipolar cells in ROs.

In conclusion, RO development recapitulated closely human retinogenesis, starting with RPCs which differentiate into the transitional populations T1, T2 and T3, followed by the appearance of all retinal neurons in a sequentially and precise order. The first-born retinal cell type were RGCs followed by horizontal/amacrine cells, cone precursors/cones, rod precursors/rods, BCPs and MCs. Remarkably, we demonstrate that the pCM region of retinal organoids may have the potential to give rise to early RPCs, suggesting their putative contribution to retinogenesis.

### Chromatin accessibility of retinal organoids is highly correlated to the developing human retina

Single cell ATAC-Seq of ROs at day 10, 20, 35, 45, 60, 90, 150 and 210 was carried out to assess chromatin accessibility during RO differentiation (**Table S5**). In total, 84,833 cells were captured using the 10XGenomics Chromium Single Cell ATAC Library and Gel Bead Kit (version 1.1). We used CellRanger to align reads and carry out data processing, and Signac/Seurat and Monocle for quality control, differential accessibility, motif enrichment and trajectory analyses. The scATAC-Seq clusters for each stage of ROs differentiation were identified using the corresponding scRNA-Seq datasets (**Fig. S7, S8**). Similarly, to scRNA-Seq analysis, defined clusters of OSE, corneal stroma, fibroblasts and corneal endothelium were easily identified: those were present mostly in the early stages of RO differentiation (up to day 60, **Fig. S7A-E**) and were much less evident at the later stages of differentiation, probably due to the incompatibility of our retinal culture media to support corneal cell survival and differentiation. Cell clusters representing optic vesicle, RPCs and NRCPs were found in the very early stages of RO differentiation (day 10 and 20) and the earliest born retinal cell types, RGCs were first identified at day 10 of RO differentiation (**Fig. S7A, B**). Notably, cell clusters of pCM, harbouring signatures of ciliary body, RPCs and pigmented cell markers were identified from day 45 of RO differentiation (**Fig. S7D** and **Table S5**), corroborating both the scRNA-Seq and the ST analyses. Notably, the three clusters of transient neurogenic progenitor cells, namely, T1, T2 and T3 were present from day 45 of RO differentiation (**Fig. S7D-E**). All the other retinal cell types emerged in an orderly fashion with horizontal and amacrine cell clusters appearing at day 35 of RO differentiation, cones from day 60, and rods, bipolar and MCs from day 90 of differentiation (**Fig. S7C-F, Fig. S8**).

Following filtering and quality control, 37,522 retinal cells from day 10-210 ROs with a 12,179 average median fragments per cell were integrated (**Fig. S8C**). We used “bedtools merge” to create a shared peak bed file, “cellranger reanalyse” to create a shared peak set with a total of 314,490 chromatin accessibility peaks, and Signac to create a gene activity matrix. Cell clustering based on chromatin accessibility peaks resulted in 17 clusters, which were identified by assaying for increases in DNA accessibility near previously known marker genes for each specific cell type (**Fig. S8C** and **Table S5**). Those included the pCM and early and late RPC clusters, as well as the T1 and T3 transient neurogenic progenitors, MCs and all retinal neurons (**Fig. S8C**). We did not identify a T2 cluster, which could be due to the transient nature of these progenitors and/or the relatively low cell number obtained for data integration following filtering and quality control.

The DNA accessibility peaks were classified using annotation from cellranger as follows: associated with promoters (if found within –1000 to +100 bp of the transcription start sites), exons, introns, distal (if found within 200 kb of the closest transcription start site), or intergenic regions (if not mapped to any genes) (**Fig. 7A** and **Table S6**). This analysis enabled the identification of cell type specific regions of accessibility for all retinal neurons arising in the ROs as well as RPCs and transient neurogenic progenitors T1 and T3 (**Fig. 7B**). We observed scATAC-Seq marker peak enrichment of cell type specific markers as follows: *WNT2B* in the pCM, *DAPL1* in the early RPCs, *FAM131C* in T1, *GAP43* in RGCs, *ONECUT1* in horizontal and amacrine cells, *OTX2* in T3, *RHO* in rods, *ARR3* in cones, *VSX1* in bipolar cells and *SLC1A3* in Muller glia cells (**Fig. 7C**).

**Figure 7:**
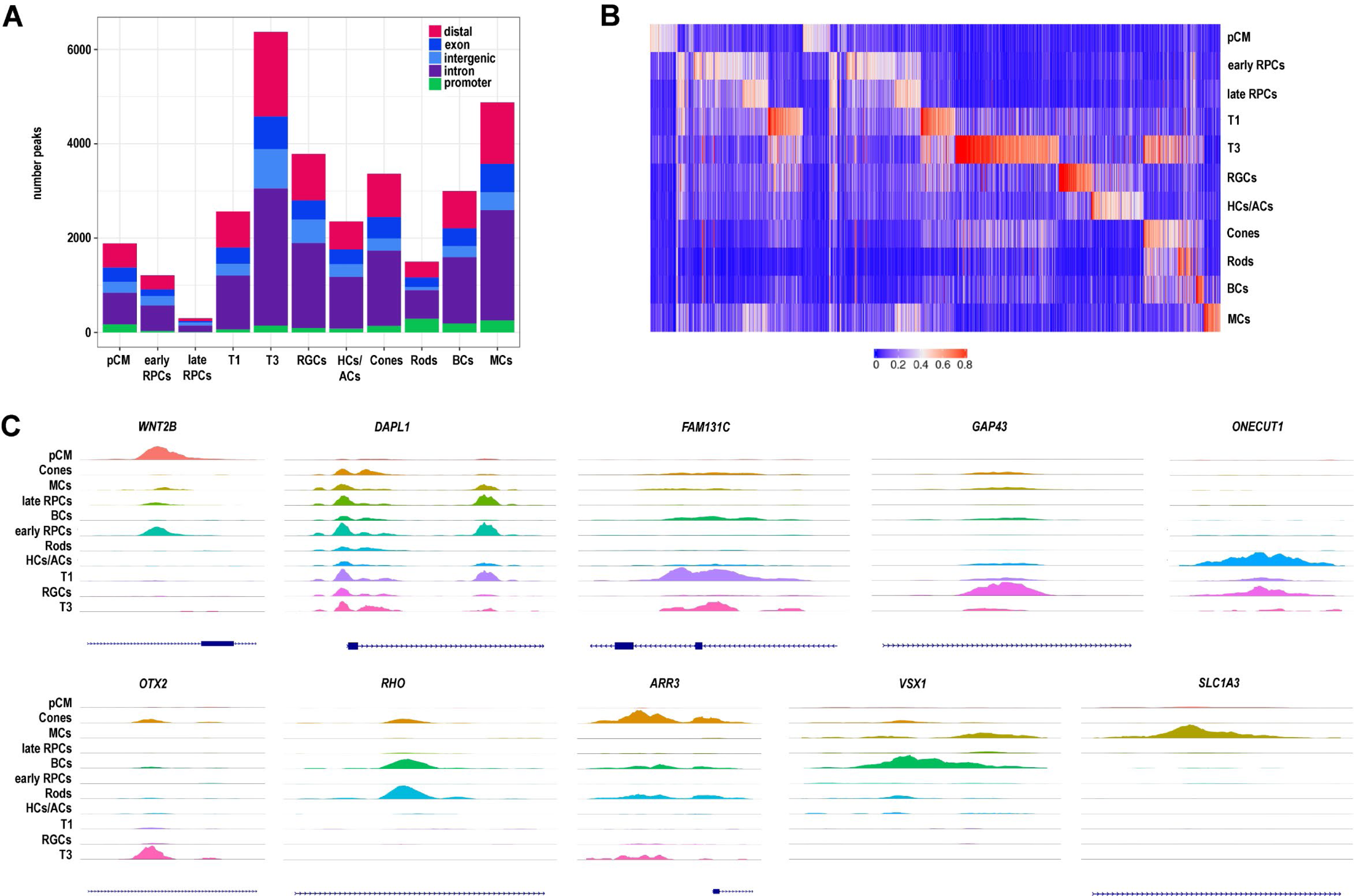
Single cell ATAC-Seq analysis of ROs indicates cell type specific chromatin accessibility profiles. **A**) The number and type of chromatin accessibility profiles for each retinal cell type found in ROs. Data are based on integrated scATAC-Seq shown in Table S5. **B**) Differentially accessible of chromatin accessibility peaks for each cell type. **C)** Representative examples of chromatin accessibility peaks for each retinal cell type individually. Each track represents the aggregate scATAC signal of all cells from the given cell type normalized by the total number of reads in TSS regions.

We identified transcription factor (TF) binding motifs using Signac, which were further validated by footprinting analysis (**Fig. 8**, **Table S7**). In the pCM, we identified binding motifs for members of the SOX, PRDM and TEAD family of TFs (**Fig. 8A, B**), corroborating the binding motifs identified by our group in RPCs of fetal retina [24]. Shared TF binding motifs were observed between RPCs and MCs for FOS/FOSL:JUN TFs, most likely reflecting the similarity in transcriptional profile between these two cell types as reported recently [17]. In accordance with generation of T3 progenitors from T1, we identified a set of shared TF binding motifs for NEUROD1, NEUROG2, HAND2 and TAL1:TCF3 (**Fig. 8A, B**). The RGCs displayed strong POU4F family member TF binding motifs, whilst horizontal and amacrine cell clusters were defined by ONECUT1, RFX and CUX TF family members binding motifs (**Fig. 8A, B**), corroborating our recent data in the developing human fetal retina [24]. As highlighted by our scRNA-Seq, cones, rods and bipolar cells are derived from the T3 neurogenic progenitors; it is therefore not surprising to observe shared binding motifs for TFs such OTX2, GSC2, PITX3, PITX2, CRX, DMBX1 (**Fig. 8A, B**), which are well described in the literature for their expression and/or role in photoreceptor specification [34–38]. Enrichment of NFI and NFX family members binding motifs was observed for MCs and BCs (**Fig. S8A, B**), consistent with their suggested role in the specification of the late-born cell types in the retina [17].

**Figure 8:**
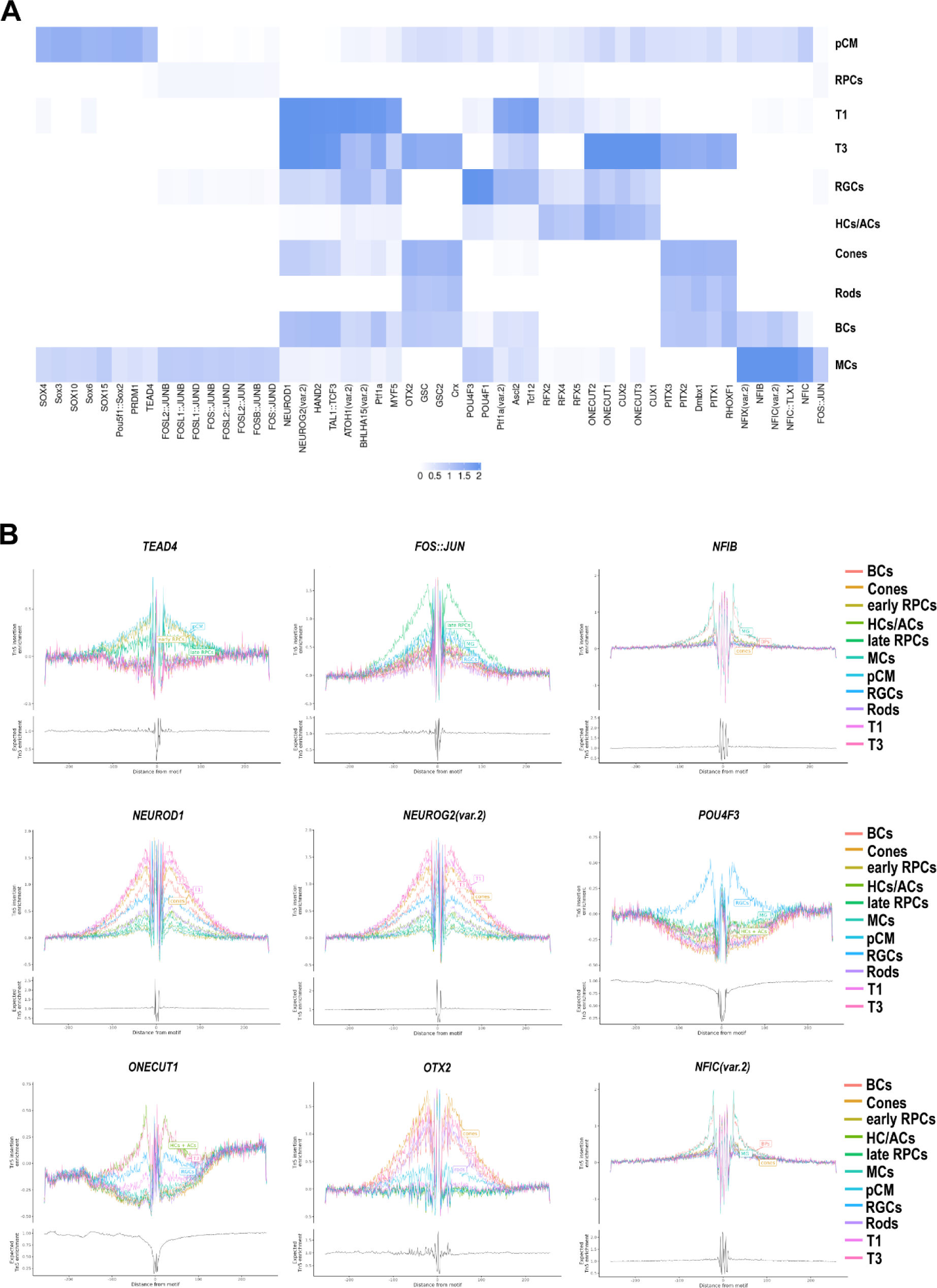
Motif analysis of accessible DNA peaks predicts cell type specific transcription factors in the retinal organoid development. **A**) Heatmap of transcription factor binding motifs enriched in each cell type. Darker colours represent more significant enrichment. **B)** Footprinting analysis of selected transcription factors predicted to display a significant enrichment in retinal cell types.

To identify gene regulatory networks that govern RPCs diversification and their differentiation, we used the Qiagen IPA upstream analysis tool combined with overlay analysis of DA peaks. In early RPCs, we predicted the activation of proliferation related upstream regulators such as MYC and MYCN, which activate expression of early RPCs (e.g., *HMGA1*) and inhibit expression of late RPCs genes (e.g., *Cyclin D1*) (**Fig. S9A**, **Table S8**). In late RPCs, several upstream regulators were identified including FGF2, TGFβ, IGF-1 and YAP1, regulating the expression of a common pool of target genes that specify late RPCs (e.g., *SOX9, FOS, VIM, CCND1;* **Fig. S9B**). Amongst the predicted inhibited upstream regulators, we found key transcription factors such as THRB and RB1 (**Table S8**) [39, 40], known for their role in the development of cone photoreceptors and retinal neurogenesis respectively (**Fig. 9C**). As RPCs differentiated to retinal neurons, we observed that the majority of predicted upstream regulators harboured an inhibitory role, perhaps demonstrating a restriction in gene expression profiles, which coupled with activation of lineage restricted transcription factors direct the differentiation to photoreceptors, RGCs and retinal interneurons. For example, CRX and NEUROD1 were predicted as upstream regulators in rod photoreceptors (**Table S8**), activating expression of key genes involved in phototransduction such as *PDE6B, PDE6A, RHO* and *SAG* (**Fig. S9D**). The major share of predicted inhibited upstream regulators was taken by miRNAs, which suppress activation of genes necessary for retinal cell fate specification. Exemplary amongst those were miR-6825-5p in cones (**Fig. S9E**) and miR-153-3p in bipolar cells (**Fig. S9F**), regulating the expression of genes encoding key proteins for cone photoreceptor genesis and function (e.g., RAX2, RXRG) and bipolar cells (e.g., NEUROD4, NFIB) respectively.

The analysis of TF binding profiles and upstream regulators showed a lot of similarities between the developing human fetal retina samples analysed by our group recently [24] and ROs studied herein. To better assess the similarity in chromatin accessibility landscape between the developing retina and ROs, we extracted the metadata from each dataset, created a unified set of overlapping differential accessibility peaks which were integrated using Harmony to remove batch effects (**Fig. S10A** and **Table S9**). Next, we ordered the human fetal retina and ROs samples in pseudotime using Monocle, revealing a nice progression from day 10 to day 210 of development with ROs being most closely situated to the equivalent age fetal retina samples (**Fig. S10B**). This was further exemplified by Spearman analysis which demonstrated 0.89 correlation between the day and pseudotime of the sample (data not shown). Pseudotime analysis for each cell type demonstrated a slightly earlier emergence of T1 neurogenic progenitors in the human developing retina samples, but similar dynamics for the T2 and T3 progenitors which emerge from T1 (**Fig. S10C**). In contrast the RGCs and cones emerged earlier in the ROs, rods and BCs later, while horizontal and amacrine cell emergence followed the timeline of human fetal retina development (**Fig. S10C**). To compare the chromatin accessibility for each cell type, we cross-correlated the pseudo-bulk DA sample in the RO to the equivalent data set obtained from our single cell ATAC-Seq analysis of developing human retina [24]. All cell types displayed a strong correlation between ROs and human fetal retina with a Spearman coefficient ranging from 0.66 and 0.69 in amacrine cells and cones respectively to 0.7 in BCs, 0.76 in rods, 0.77 in horizontal cells, 0.79 in MCs and 0.89 in RGCs (data not shown). Together these data demonstrate that the chromatin accessibility of ROs is highly correlated to the developing human retina, but with some differences in the temporal emergence and abundance of the retinal progenitors and mature cell types.

## Discussion

ROs generated from human PSCs provide a tractable platform for studying early retinal development and disease. Several recent studies have used scRNA and/or ATAC-Seq to reveal cell type composition in ROs and to identify cell specific cis-regulatory elements and gene regulatory networks [10, 12, 14, 15, 22, 41–43]. Recently Wahle and colleagues (2023) developed a multiplex immunohistochemistry approach to establish a spatial retinal organoid map of retinal cells during development, but this was based only on expression of 63 antibodies covering the majority of the retinal cell types [44]. Herein, using a combination of scRNA-Seq and ST approaches, we present for the first time a genome wide, single cell spatio-temporal transcriptome map of RO development, revealing the sequential and precise order of all retinal neuron emergence and their specific localisation, which closely recapitulates human retinogenesis. Importantly, we provide evidence for the existence of a pCM domain at the very edge of ROs, from where a population of early RPCs are likely to arise. Combining the scRNA-with scATAC-Seq data, we were able to reveal cell type specific TF binding motifs on accessible chromatin at each stage of RO development and to show that chromatin accessibility of ROs is highly correlated to the developing human retina, but with some differences in the temporal emergence and abundance of some of the retinal cell types.

The human retinal ontogenesis starts very early during human development. At 2 weeks post conception the optic vesicle evaginates to form the optic cup, whose inner layer differentiates into neural retina [45]. These processes represent fundamental events for retinal ontogenesis, however due to scarcity of available human retinas of this age, most of our molecular and developmental data so far have been acquired from animal models. To fill this gap in our understanding of retinal development, we have used very early ROs (day 10 and 20), revealing for the first time the molecular signature of optic vesicle and optic cup neuroepithelium at the single cell level, and most importantly their spatial localisation, thus providing novel information and early molecular markers for assessing retinal development *in vitro* and *in vivo*. We were able to detect a large pool of RPCs at day 20 of RO differentiation using both scRNA– and ATAC-Seq. ST revealed that the RPCs were initially located all over the ROs, but from day 35 the RPCs localised at the apical edge of the organoids, whilst the NRPCs were predominantly found on the basal side of the organoids. Notably from day 45, we were able to identify with all three methods a domain located at the very edge of ROs with transcriptional signatures of RPCs, ciliary body and pigmented epithelium, which we defined as a pCM based on similar transcriptional signature we revealed by spatiotemporal single cell analyses of developing human retina in a recent study [24]. A similar entity was described by Kuwahara and colleagues (2015) who used IF imaging and pulse labelling studies to show that the pCM region can expand the neural retina by *de novo* progenitor generation [2]. In accordance with this, our pseudotime analysis positions the pCM at the start of RPCs pseudotime indicating that some of the earliest RPCs in the organoids are likely to emerge from this region. If this were to be the case, the question remains of why this domain was not detected at the earlier stages of organoid development? Could this be due to two separate sources of RPCs: one directly arising from differentiation of optic cup to retinal neuroepithelium and the second from the pCM domain which is established slightly later in development? Or could this be due to the small size of pCM in earlier organoids? To answer this question, lineage tracing combined with single cell RNA-Seq and ST need to be performed as recently demonstrated for first heart field predominance of human PSC differentiation [46].

RGCs developed in ROs have been reported to be progressively lost in long-term cultures [5], probably due to demand for additional reagents in the culture media that promote their survival. In contrast we were able to detect RGC clusters from day 10 and day 20 until the end of the differentiation time point (day 210) by both scATAC– and scRNA-Seq methods respectively and able to locate them predominantly on the basal side of ROs using ST. This could be due to the presence of IGF-1 in our culture media, which has been shown to enhance RGC survival *in vivo* [47]. Strikingly, we were able to detect dying RGCs were in the centre of ROs, likely reflecting the drastic reduction of RGCs during human retinal development. The similarity of RGCs to their *in vivo* counterparts in the developing human retina was further demonstrated by a very strong correlation (0.89) revealed by the comparison of the accessible chromatin peaks.

Horizontal and amacrine cell clusters were identified by both scRNA– and ATAC-Seq methods from day 60 and day 35 of RO differentiation respectively, but not always as separate clusters, which could be due to the high similarity in their transcriptional and chromatin accessibility profiles and/or low abundance in organoids. This was also the case for the transient neurogenic progenitor population T2, which give rise to both of these cell types, but was often difficult to be identified as a separate cluster in the later stages of RO differentiation (day 150 and day 210). Comparison of abundance of these cell types using the chromatin accessibility and transcriptional data demonstrated a much lower percentage of horizontal, amacrine and T2 progenitors in the ROs compared to human fetal retina (data not shown), suggesting that the 3D environment and/or culture conditions are not fully supportive of the development of these progenitors/interneurons. These findings corroborate published evidence showing a disorganisation of the inner layer of organoids at approximately day 70 to day 90 of development [22].

Our data demonstrate that cones arise slightly earlier and rods slightly later in ROs, compared to their emergence *in vivo*. Both types of photoreceptors were localised by ST in the apical layer of ROs, with rods located close to the brush border of ROs and cones just underneath those. BCs and MCs were the last cell types to develop in ROs, corroborating previously published data for ROs and fetal retina [10, 17, 33]. BCs developed slightly later in ROs compared to human fetal retina and demonstrated a high transcriptional similarity with T3 progenitors, which led us to suggest that these more likely represented BCPs, rather than mature BCs.

Recently Fernando and colleagues (2022) described the formation of brain and ROs from pluripotent stem cells connected by nerve-like axonal projections of optic origin using a 2D-3D method [48]. While such organoids were not detected in our culture system, another type of organoid which contained a retina-like core, but also an opaque cystic-like structure within which corneal epithelium, stroma, endothelium and lens cell clusters was detected, corroborating previously published data by our group [49]. Given the presence of corneal, retinal and lens cells within the same organoid, we refer to those as “eye-like organoids.” These organoids did not survive beyond day 90 of differentiation, suggesting that an adapted media composition is necessary to further promote their survival and maturation.

The presence of astrocytes in the retina was first described by Cajal [50], however their origin has remained under discussion for a long time, with some proposing the astrocytes to be derived from transformed MCs and others suggesting that these cells may arise *in situ* from the retinal neuroepithelium [51] as they do in spinal cord grafts in avians [52]. Strikingly, ST revealed the presence of a small astrocyte cluster in day 210 ROs, characterised by high expression of GFAP and VIM, which may suggest that these could represent immature astrocytes [53]. In rodents, astrocytic cells migrate into the retina through the optic nerve head and then spread towards the periphery across the nerve fibre layer [53]. Moreover, the optic nerve serves a as the source of astrocyte progenitors, thus the presence of the optic stalk is critical for astrocyte development. We did not detect cell clusters with transcriptional signatures of optic stalk at day 210 of differentiation; thus, we believe that origin of astrocytes in humans may differ from rodents, however this remains to be further investigated. Our findings echo recent published data showing the presence of donor derived astrocytes into retinas of mice transplanted with micro-dissected multi-layered retinal fragments from human ROs [54]. It not clear if astrocyte progenitors were present in the ROs in small numbers or transplantation induced trans-differentiation of neural retinal cells into astrocytes. However, given the presence of astrocytes in ROs studied herein, we are tempted to speculate that astrocyte progenitors do arise in ROs, although their cell of origin remains as yet unknown.

Application of all three techniques on ROs samples of the same age enabled us to compare and contrast our results. Similarly, to our findings in human fetal retina [24] we observed that scATAC-seq preceded cell date identification compared to scRNA-Seq. For example, a small RGCs cluster was identified at day 10 ROs in the scATAC-Seq, but only in day 20 by scRNA-Seq. Likewise, BCs and MCs were found at day 90 in scATAC-Seq while scRNA-Seq indicated their emergence at day 150. This suggests that the chromatin accessibility changes, and TF binding occur in advance of cell specific gene expression changes that drive retinogenesis. We also noticed that rare clusters were not always identified by ST. These included the transient neurogenic progenitors and interneurons, which often were detected as mixed clusters of horizontal, amacrine cells and RGCs, probably due to technical limitations of the ST slides.

## Study limitations

Due to relatively high cost of single cell analyses, all the results presented in this study are derived from ROs generated from a single human iPSC (hiPSC) line, with validations performed in ROs generated from one human embryonic stem cell (hESC) line. It would be useful to expand all the three single cell analyses on ROs generated from additional PSCs grown under the same as well as different culture conditions. We also used the Visium ST system which has 4992 barcoded gene expression spots per capture area where each spot has diameter of 55µm with a 100µm centre to centre distance between spots. This was insufficient in our experience to define all the progenitor and retinal cell types. It is thus desirable to use a higher definition ST method and combine it with protein detection to better define rarer cell types and localise them within the ROs. Cell barcoding combined with single cell analyses are currently being tested in a few model systems [46, 55]: these methods combined would provide better insights into sources of RPCs and their differentiation trajectory. Despite these minor limitations, our work provides the scientific community with a powerful spatiotemporal single cell atlas of human PSCs undergoing differentiation to laminated ROs.

## Methods

### Retinal organoid differentiation

hiPSCs (WT2) [56, 57] were expanded in mTESR™1 (StemCell Technologies, 05850) on growth factor reduced Matrigel coated plates (BD Biosciences, San Jose, CA) at 37°C and 5% CO_2_. For the generation of ROs, confluent hiPSCs were dissociated into single cells using Accutase (Gibco, A1110501) and were seeded at a density of 7,000 cells/well onto Lipidure (AMSbio, AMS.52000011GB1G) pre-coated 96-well plates (U-bottom, Helena, 92697T) in mTeSR™1 supplemented with 10 μM Y-27632 ROCK inhibitor (Chemdea, CD0141). 200 μl of differentiation medium was added after 2 days and hereupon half of the differentiation medium was changed every 2 days until day 18 of differentiation as described in Dorgau et al. (2018) [58]. After day 18, the differentiation media was supplemented with 10% Fetal Calf Serum (FCS; Thermo Fisher), Taurine (Merck) and T3 (Merck) and ROs were further cultured in 6-well low attachment plates (Corning, 3471). Retinoic Acid (RA; 0.5 µM, Merck) was added from day 90 to day 120 of differentiation. The media was changed every 2-3 days for remaining differentiation.

Prior to ROs collection, brightfield images were taken using a AxioVert upright microscope (Zeiss, Germany). ROs were collected at different developmental stages for experiments, starting with early developmental samples (day 10, 20 and 35), followed by the mid developmental (day 45, 60 and 90) and late developmental (day150 and 210) samples.

### scRNA– and –ATAC-Seq

ROs samples of different developmental stages (day 10, 20, 35, 45, 60, 90, 150 and 210) were dissociated to single cells using a neurosphere dissociation kit (Miltenyi Biotech) (**Tables S1, S3, S5**). 10,000 cells from each sample were captured, and sequencing libraries generated using the Chromium Single Cell 3’ Library & Gel Bead Kit (version 3.1, 10x Genomics) for scRNA-Seq. For scATAC-Seq 10,000 of the subsequent nuclei were captured, and sequencing libraries generated using the Chromium Single Cell ATAC Library & Gel Bead Kit (version 1.1, 10x Genomics). Single cell RNA-Seq libraries were sequenced to 50,000 reads per cell and scATAC-Seq libraries were sequenced to 25,000 reads per nucleus on an Illumina NovaSeq 6000.

### scRNA-Seq analysis

Cellranger mkfastq version 3.01 was used to de-multiplex the BCL files into FASTQ files. Reads were then aligned and quantified using Cellranger count and the human reference genome GRCh38. The quality of the cells in each sample were checked in R. Cells which had fewer than 1000 reads or 500 genes or more than 10% mitochondrial reads were removed from downstream analysis. DoubletFinder was used to identify doublets within the data these were then removed from the datasets. Each sample was assessed individually following the Seurat (version 4.3.0). The standard clustering workflow was followed. This included normalisation to remove cell to cell differences, scaling the data to remove differences in expression levels between genes, identification of a set of 2000 biologically informative highly variable genes and principle component reduction of data, and clustering of the data. The following factors were regressed out during the scaling process, “percent.mt”, “nCount_RNA”, “nFeature_RNA”.

Retinal cells were then selected from the individual samples and batch effects removed by Harmony (version 0.1.1) to create an integrated dataset. The data was visualised using a Uniform Manifold Approximation and Projection (UMAP) based on the first 10 batch corrected coordinates and clusters identified using Seurat. Differentially expressed markers between each cluster were identified using the Seurat FindMarkers function with the method Wilcoxon test. These genes were used to identify the cell types in the clusters shown in **Table S1 and S3**.

The combined dataset was subset to study development trajectory within the following cell types: pCM-RPC-T1-T2-T3, pCM-RPC-T1-T2-HC-AC-RGC, pCM-RPC-T1-T3-Cones-Rods-BC (**Table S4**). Cells within each of these groups were selected and then the data was re-clustered using the method described for the full dataset. We then reassessed the annotations to ensure that we had high confidence in the cell type assignment. Pseudotime trajectories for each developmental branch were created using Monocle 3. Genes which were differentially expressed between cell types were identified within the subsets. These were visualised in heatmaps where cells were ordered within cell type by pseudotime.

### scATAC-ATAC analysis

Cellranger ATAC software (version 1.2) was used to identify peaks in each of the samples. Bedtools merge (version 2.30) was used to create shared peaks and the data was reanalysed using Cellranger ATAC reanalyse. Quality control steps were performed using Signac. The datasets were imported using Signac and quality control steps were performed to remove cells low quality cells. We filtered cells which had fewer than 20% of reads in peak region fragments, or less than 3000 peak region fragments. Cells with a TSS enrichment score of less than 2 and Blacklist ration greater than 0.05 or a nucleosome signal of less than 4 were also removed from downstream analysis.

The standard Signac workflow to cluster the cells, which consists of term frequency-inverse document frequency (TF-IDF) normalisation; singular value decomposition (SVD) dimension reduction; followed by UMAP reduction was applied to the data. Seurat was then used to cluster the cells. LSI components 2 to 30 were used to create the UMAP and for clustering analysis. A gene activity matrix was generated using Signac. This was used to predict upregulated genes in each cluster and assign the clusters to cell types (**Table S5**).

A combined dataset was created by selecting retinal cells from each sample. Retinal cell types were selected from each sample and a combined dataset was created using the method described in the previous paragraph. The Logistic regression (LR) test from the FindAllMarkers function was used to identify differentially accessible peaks for each of the annotated cell types (**Table S6**). The ComplexHeatmap package was used to plot the average peak value for each differentially accessible peak for each cell type.

The per cell motif activity was computed using Chromvar. Enriched motifs for each cell type were identified using FindAllMarkers. ComplexHeatmap was used to show the top cell type enriched motifs ordered by average difference in z-score (**Table S7**). Signac was used to generate motif plots. Qiagen Ingenuity Pathway Analysis (IPA) was used to analyse differentially accessible peak lists and identify upstream regulators. We looked for a consensus in the analysis by overlaying the predicted motifs onto the IPA promoter predictions (**Table S8**).

### ST

RO samples were collected and fresh frozen at day 10, 20, 35, 45, 60, 90, 150 and 210 of differentiation (**Table S2**). Each sample contained 15-25 retinal organoids, depending on the stage of differentiation. Sections (10µm) were cut (Leica Cm1860) and carefully placed into the capture areas of the spatial transcriptomic slides. Slides were stored at –80°C until needed. Spatial transcriptome analyses were performed with the Visium Spatial Gene Expression kit from 10XGenomics. First, the tissue optimisation was performed defining 24-30 minutes as the most optimal permeabilization time window. The gene expression ST procedure was performed according to manufacturer’s instructions. Briefly, sections in the four capture areas for each sample were fixed, haematoxylin and eosin (H&E) stained. To preserve histological information H&E-stained sections were imaged using a NikonTiE inverted microscope at 10x magnification (Plan Fluor 10x 0.3 NA). Image tiles captured and stitched together using Nikon Elements software, enabling to overlay the cell tissue image and the gene expression data later. After permeabilization, reverse transcription reagents were added on top of the tissue sections. The tissue sections were subsequently removed, leaving the cDNA coupled to the arrayed oligonucleotides on the slide. Then the cDNA-RNA hybrids were cleaved off the chip and the sequencing libraries were prepared. The sequencing depth varied between 100,000,000 and 250,000,000 million reads.

### ST analysis

The data was de-multiplexed, aligned and quantified using Spaceranger version 1.0. The data was aligned to the human reference genome GRCh38. Spaceranger generated spot co-ordinates calculated using fiducial detection, and regions under the tissue section. Spaniel (version 1.12) was used to import the data into R. The 4 consecutive sections tissue sections from each sample were clustered using the Seurat scRNA-Seq pipeline described above. A resolution of 0.5 was used for the cluster analysis. Differentially expressed genes were identified between clusters. The spatial expression and cluster plots were generated using Spaniel.

### RNAscope

RNAscope *in situ* hybridization assay was used to determine the expression profile of *ZIC1* and *HES6* during human retinal organoid development. Fresh frozen sections of ROs aged day 45 as used for spatial transcriptomic experiments were formalin fixed for 1 h at 4°C. The sections were dehydrated through a series of ethanol 50%, 70% and two changes of 100%. The sections were incubated with a protease IV (ACD-Cat. No. 322336) for 15 min at 40°C. RNAscope probes Hs-ZIC1-C2 (ACD-Cat No. 542991-C2) and Hs-HES6-C3 (ACD-Cat No. 521301-C3) were hybridised to the tissue for 2 h at 40°C followed by multiple rounds of signal amplification. Positive (ACD-Cat No. 320861) and negative (ACD-Cat No. 320871) control probes were used to confirm specificity. The annealed probes were detected using Opal fluorophores OPAL 650 (C2) and OPAL 520 (C3) and imaged using an Axio Imager upright microscope with Apotome structured illumination fluorescence and the ZEN software.

### IF analyses

‘Eye-like’ ROs were collected at day 45 and day 90 of differentiation. After fixation using 4% paraformaldehyde (PFA; Santa Cruz Biotechnologies) organoids were cryopreserved, embedded in OCT (Cell Path Ltd, Newtown, UK) and cut (10 µm) on a cryostat (Leica Cm1860). IF was performed as previously described [59]. Sections were reacted against the following primary antibodies: CRYAA (1:50; gift from Roy Quinlan), CRYAB (1:500, Abcam, ab76467), CRX (1:200, Abnova, H00001406-MO2), Ki67 (1:200, Abcam, ab15580), KRT13 (1:500, Abcam 52816), KRT15 (1:500, Abcam, ab92551), Lumican (1:50, Santa Cruz Biotechnologies, sc-166871), MT2A (1:100, Sigma-Aldrich, SAB1402848) and NP63 (1:250, Abcam, ab735). Secondary antibodies were conjugated to Alexa488 (Jackson Immuno Research Laboratories) and Cy3 (Jackson Immuno Research Laboratories). Antibody specificity was assessed by omitting the primary antibodies. Images were obtained using a Zeiss Axio Imager.Z1 microscope with ApoTome.2 accessory equipment and Zen software. Representative images are displayed as a maximum projection and adjusted for brightness and contrast in Adobe Photoshop CS6 (Adobe Systems).

### Comparison of organoid samples with human retina

All organoid ATAC-seq samples were compared with 12 ATAC-seq samples of developing human eyes and neural retina from 10-21 post-conception weeks (GSE234003) [24]. A unified set of overlapping peak ranges from both datasets were created. Peaks were filtered if they were greater than 10000bp or less than 20bp. Cellranger-atac was used to call peaks in all organoid and human retina samples using the shared ranges. The data was then processed following the Signac workflow described above. Harmony was used to batch differences between the human fetal retina and organoid samples. The data was clustered at a resolution of 0.8 and 36 clusters were identified and visualised on a UMAP plot. These were annotated as cell types using the gene activity values (**Table S9**). Monocle 3 was then used to create a pseudotime trajectory. The median pseudotime values for each sample were then plotted against age in days. Beeswarm plots were used to compare the pseudotime range of organoid and human cells for each cell type.

## Data availability

All the single cell data have been deposited to GEO under the following accession numbers: GSE235577.

To review GEO accession GSE235585

## Acknowledgements

The authors are grateful for funding received from BBSRC UK (BB/T004460/1), MRC UK (MR/S035826/1, MR/S036237/1). The authors would like to acknowledge the Bioimaging Unit (Newcastle University) for their support and assistance in this work and Prof. Roy Quinlan (University of Durham) for the kind gift of CRYAA antibody. This publication is part of the Human Cell Atlas – www.humancellatlas.org/publications/.

## Author contributions

BD –performed experiments, data collection and analyses, figure preparation and manuscript writing

JC – performed experiments, data collection, fund raising

AR, VB, MMM, RH, JC, TD, LA – performed experiments, data collection

RQ – bioinformatic data analyses, data submission, figure preparation, contributed to manuscript writing and fund raising

ML – experimental design, data analyses, figure preparation, manuscript writing and fund raising

