## Supplement Figure S1-10, Table S1-9 for "Deciphering the spatio-temporal transcriptional and chromatin accessibility of human retinal organoid development at the single cell level"

**This PDF file includes:**

Supplementary Text
Figs. S1 to S10
Tables S1 to S9

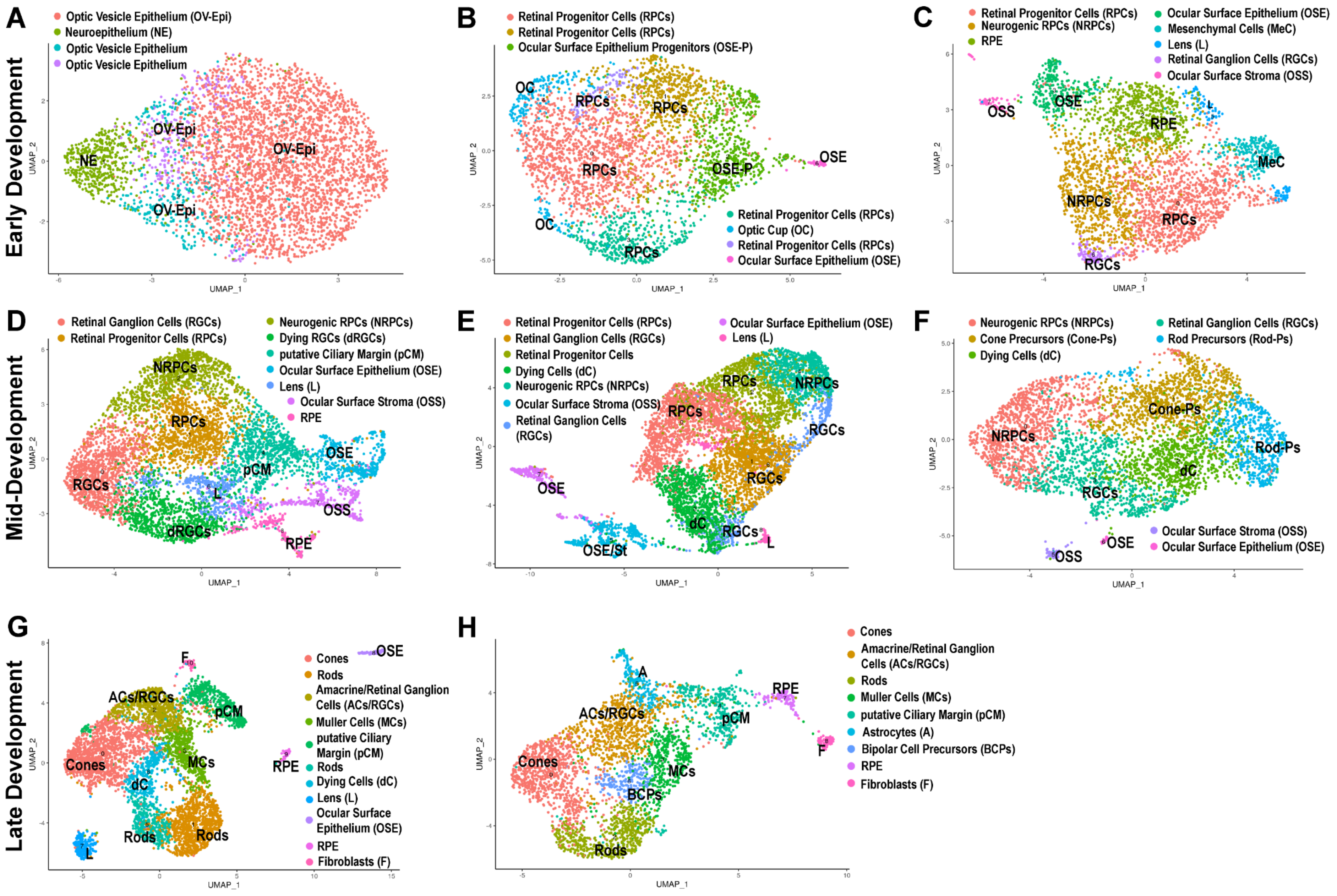


**Figure S1: Gene clusters identified in ST analyses in early, mid and late retinal development. A-H)** UMAP plots showing individual stages during early RO differentiation, including day 10 (**A**), day 20 (**B**) and day 35 (**C**), mid retinal development, composed of day 45 (**D**), day 60 (**E**) and day 90 (**F**) and late retinal development with day 150 (**G**) and day 210 (**H**). Cluster annotations are based on highly expressed retina specific marker genes, shown in Table S2.

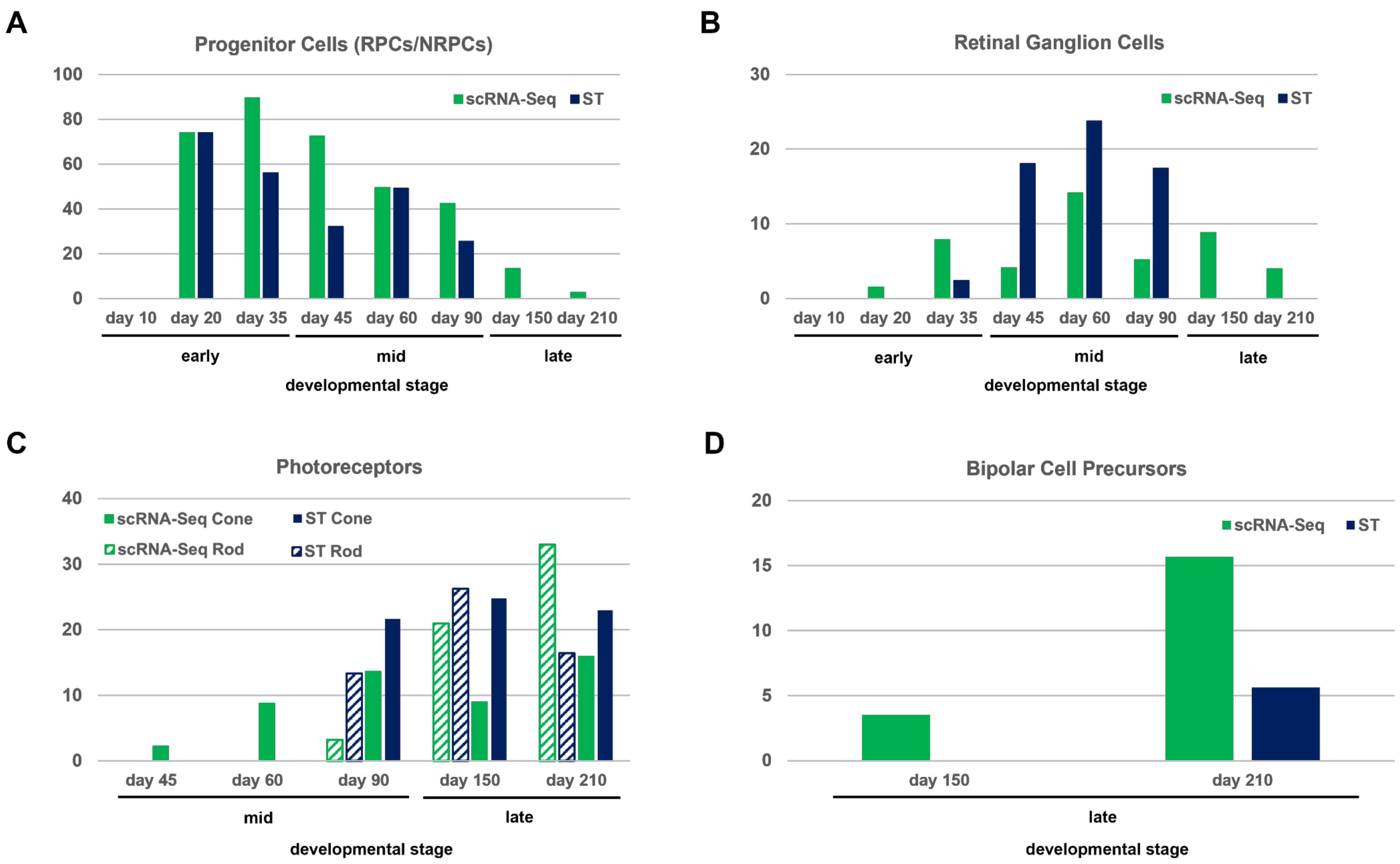


**Figure S2: ScRNA-Seq and SST analyses comparison of retinal cell type development during RO differentiation. A)** Progenitor cell population (RPCs/NRPCs) peaks at day 20 in ST data and day 35 in scRNA-Seq data and then decreases over time. **B)** RGCs population increases up to day 60 of differentiation and diminishes afterwards. Note RGCs were excluded at day 150 and 210 differentiation from ST analyses as they were detected in a mixed cluster with amacrine cells. **C)** Photoreceptors (cones and rods) expand from day 45 onwards during RO development with both techniques, even so cones emerge earlier than rods. **D)** BCPs emerge from day 150 onwards in scRNA-Seq analysis, revealing a population increase until day 210: those were first detected at day 210 of differentiation in ST analysis.

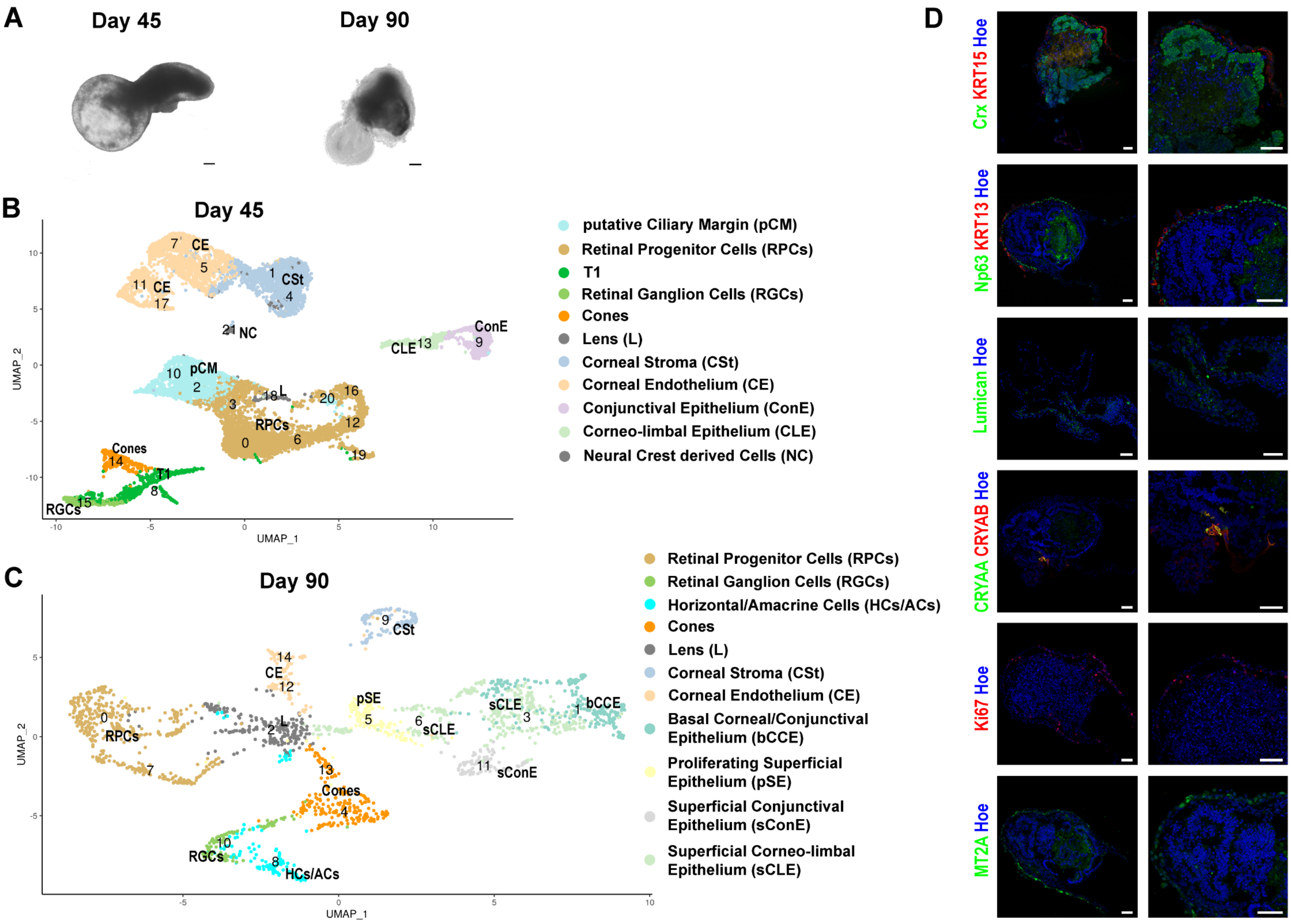

A
Day 45
Day 90
B
Day 45
UMAP_2
UMAP_1
CE
CSt
NC
ConE
CLE
pCM
L
RPCs
Cones
T1
RGCs
putative Ciliary Margin (pCM)
Retinal Progenitor Cells (RPCs)
T1
Retinal Ganglion Cells (RGCs)
Cones
Lens (L)
Corneal Stroma (CSt)
Corneal Endothelium (CE)
Conjunctival Epithelium (ConE)
Corneo-limbal Epithelium (CLE)
Neural Crest derived Cells (NC)
C
Day 90
UMAP_2
UMAP_1
CSt
CE
pSE
sCLE
bCCE
sConE
RPCs
L
Cones
RGCs
HCs/ACs
Retinal Progenitor Cells (RPCs)
Retinal Ganglion Cells (RGCs)
Horizontal/Amacrine Cells (HCs/ACs)
Cones
Lens (L)
Corneal Stroma (CSt)
Corneal Endothelium (CE)
Basal Corneal/Conjunctival Epithelium (bCCE)
Proliferating Superficial Epithelium (pSE)
Superficial Conjunctival Epithelium (sConE)
Superficial Corneo-limbal Epithelium (sCLE)
D
Crx KRT15 Hoe
Np63 KRT13 Hoe
Lumican Hoe
CRYAA CRYAB Hoe
Ki67 Hoe
MT2A Hoe

**Figure S3: Emergence of eye-like organoids at the mid stages of organoid differentiation. A)** Morphology of eye-like organoids at day 45 and day 90. **B, C)** scRNA-Seq UMAP plots of eye-like organoid cells at day 45 (**B**) and day 90 (**C**) revealing a retinal core with RPCs, T1 transient neurogenic progenitors, RGCs and cones alongside lens and cornea which displays the formation of all layers: the endothelium, the stroma and the epithelium. Annotation of each cluster is based on highly expressed markers, shown in Table S3. **D)** IF analysis of eye-like organoidss at day 45 of differentiation, confirming the presence of a retinal core by Crx-positive cells (green) together with the presence of lens (CRYAA, green and CRYAB, red), limbal-corneal epithelium (KRT15, red), conjunctival epithelium (KRT13, red), corneal stroma cells (Lumican, green) and proliferative epithelial progenitor cells, identified by the expression of Np63 (green), MT2A (green) and Ki67 (red). Cell nuclei were counterstained with Hoechst. Scale bars, 20 µm.

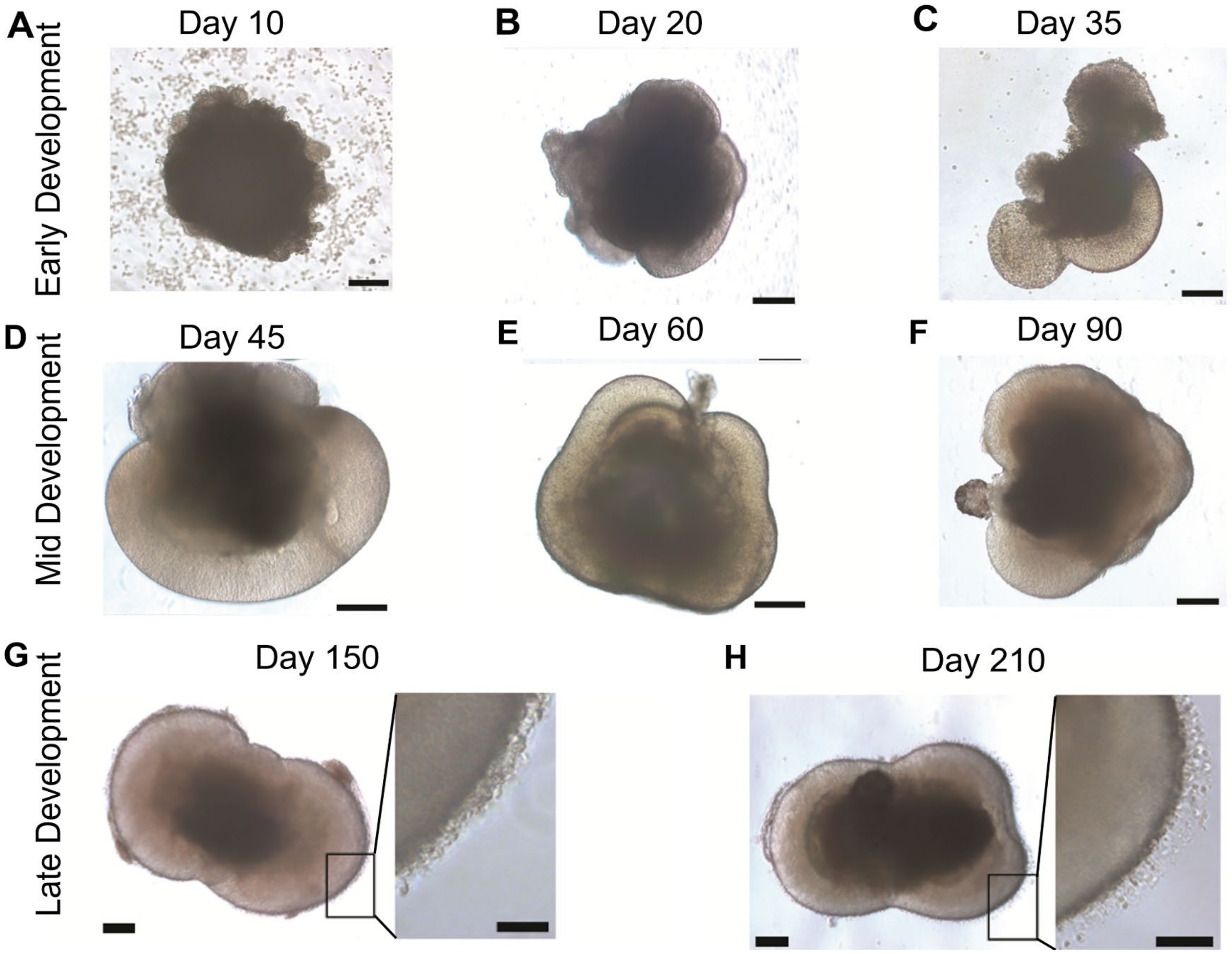


**Figure S4: Morphological development of ROs during the course of differentiation. A-C)** Representative brightfield images of ROs at day 10 (**A**), day 20 (**B**) and day 35 (**C**) during early differentiation, revealing the formation of neuroepithelium at the apical edge of ROs from day 20 (**B**) onwards. **D-F)** ROs in mid developmental stages, day 45 (**D**), day 60 (**E**) and day 90 (**F**) of differentiation indicate the thickening of the neuroepithelium at the apical edge of organoids. **G, H)** ROs at day 150 (**G**) and day 210 (**H**) mature during late development, forming inner/outer segments ("brushborder", higher magnifications) at the apical edge of organoids. Scale bars, 100 µm and 50 µm for higher magnifications.

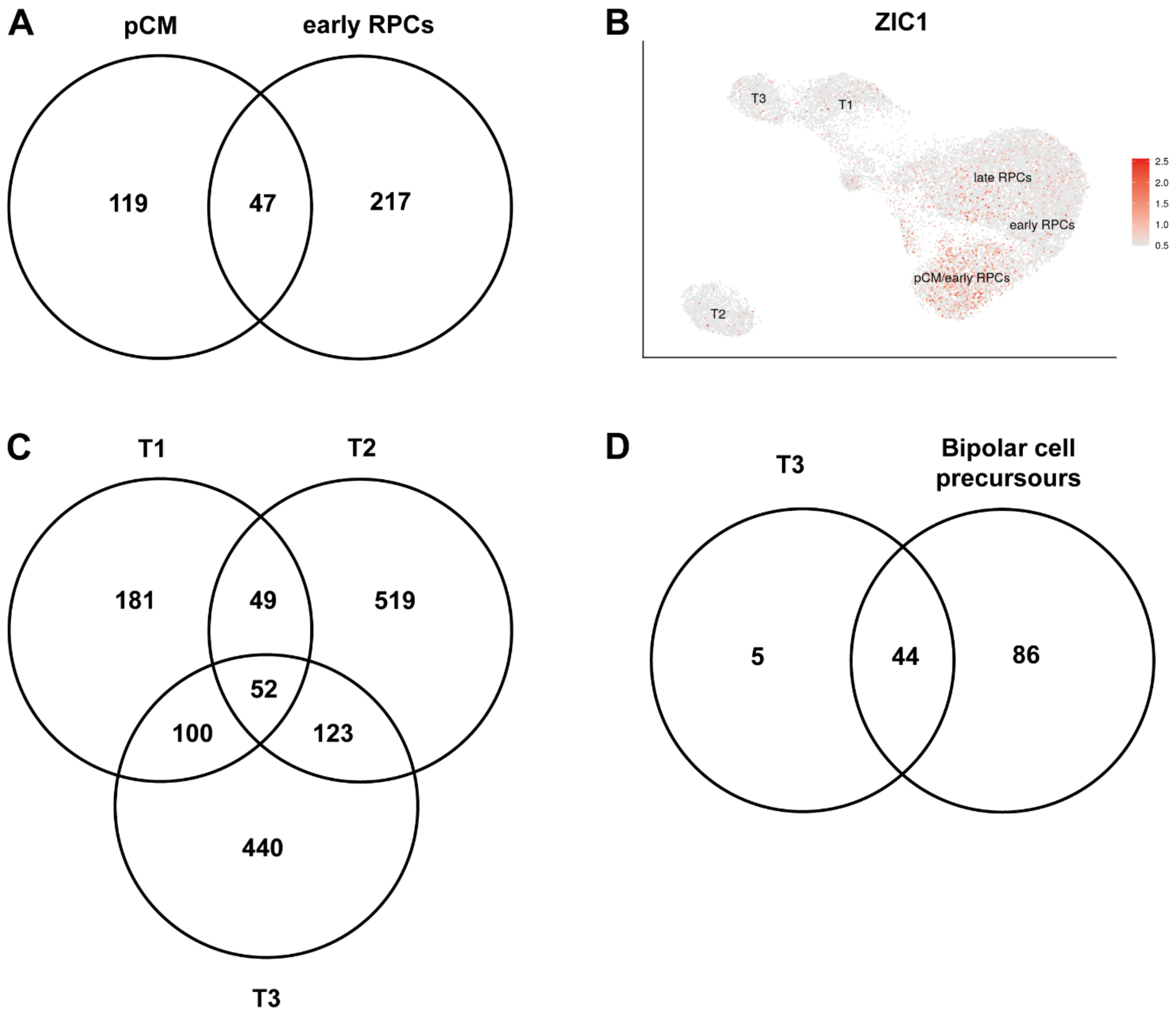


**Figure S5: Gene expression profile comparison between the pCM and early RPCs, neurogenic transient populations (T1-3) and neurogenic transient population T3 with bipolar cells indicate overlapping as well as unique marker genes for cell type. A)** Overlapping gene expression pattern of pCM and early RPCs, indicating some early RPC-like cells in the pCM zone of ROs. **B)** *ZIC1* expression profile in the pCM, RPCs, and transient neurogenic progenitors (T1-T3) pseudotime trajectory showed highest expression in the pCM of ROs. **C)** Gene expression comparison between transient neurogenic progenitors, T1 - T3, revealing some similarities between all clusters. **D)** BCPS indicated a high transcriptional similarity to transient neurogenic progenitor population, T3, suggesting the lack of mature BCs in ROs.

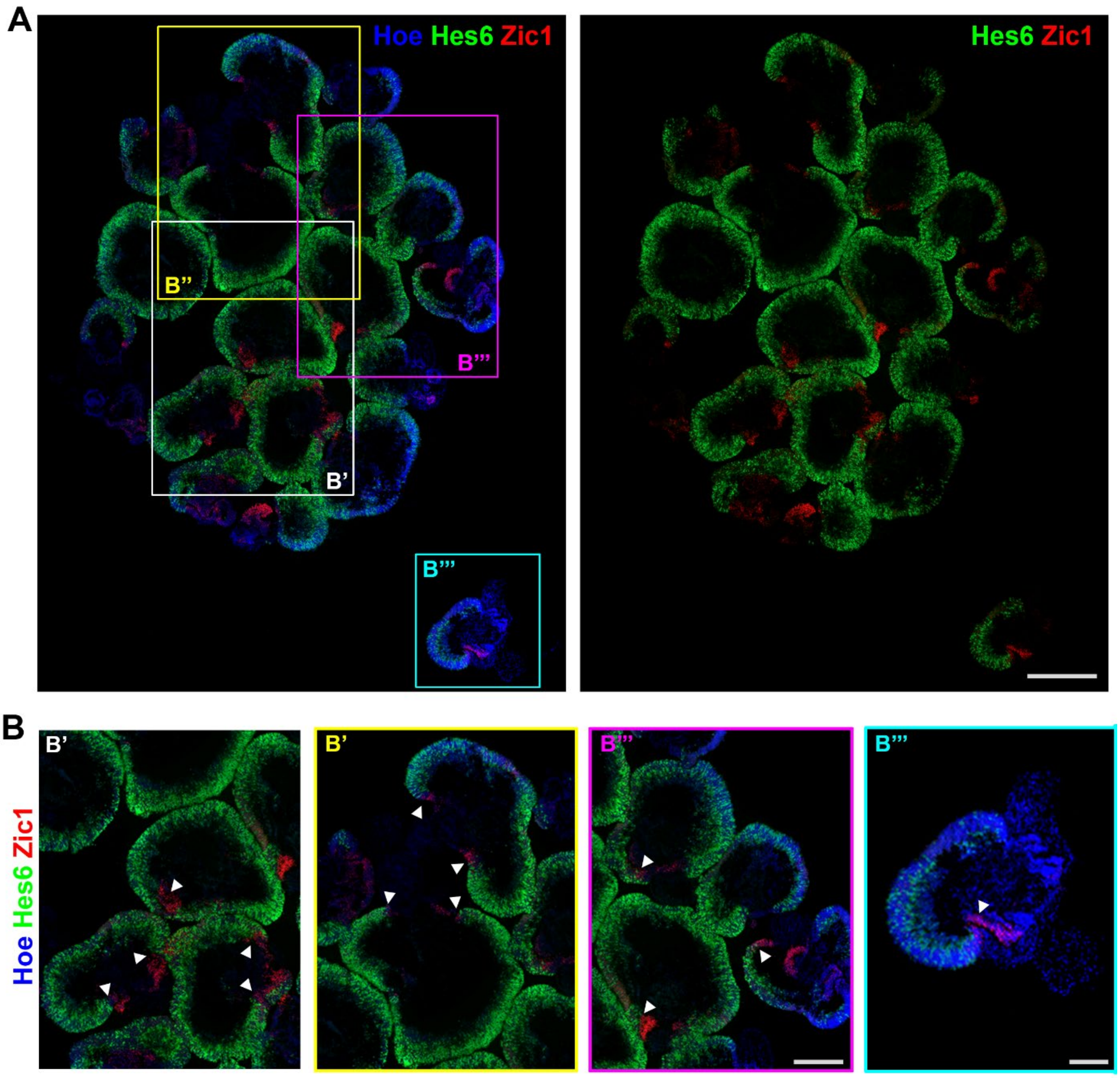


**Figure S6: RNAscope reveals strong expression of early RPC marker *ZIC1*, in the pCM at day 45 of differentiation, while the late neurogenic RPC marker HES6 expression marks the rest of retinal neuroepithelium in a complementary pattern to *ZIC1*. A)** Expression of *HES6* (green) and *ZIC1* (red) in ROs at day 45 of differentiation, confirming restricted expression of ZIC1 in pCM. **B)** Higher magnification images showed the restricted *ZIC1* expression in the pCM of ROs and its spatial location at the distal tip of the retinal-like retinal structure (white arrowheads). Cell nuclei are counterstained Hoechst (Hoe, blue). Scale bars, 50 µm for A and 10 µm for B**.**

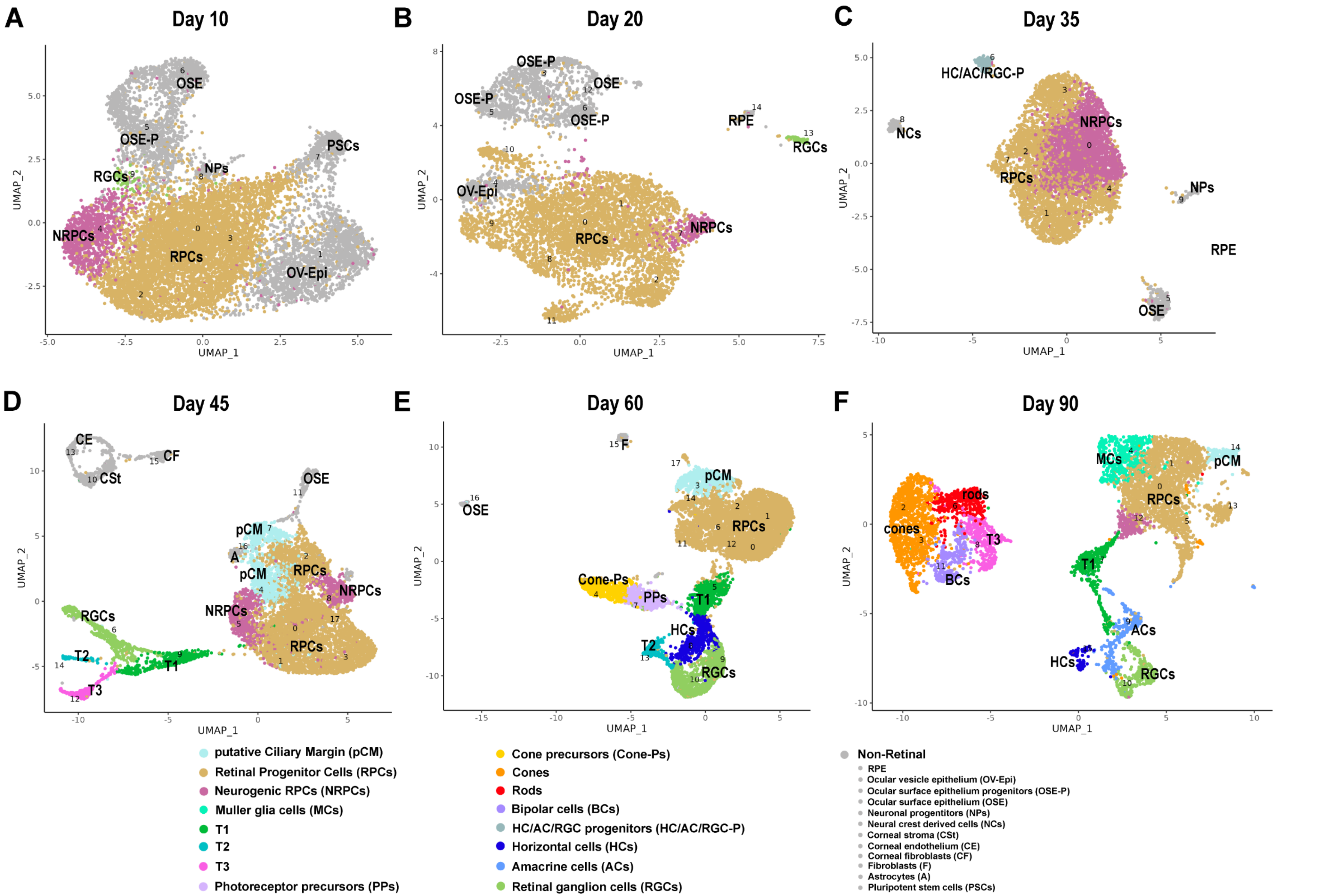
A
Day 10
OSE
OSE-P
NPs
PSCs
RGCs
NRPCs
RPCs
OV-Epi
B
Day 20
OSE-P
OSE
RPE
RGCs
OV-Epi
RPCs
NRPCs
C
Day 35
HC/AC/RGC-P
NCs
NRPCs
RPCs
NPs
RPE
OSE
D
Day 45
CE
CF
CSt
OSE
pCM
A
RPCs
NRPCs
RGCs
T2
T1
T3
E
Day 60
F
OSE
pCM
RPCs
Cone-Ps
PPs
T1
HCs
T2
RGCs
F
Day 90
MCs
pCM
RPCs
rods
cones
T3
BCs
T1
ACs
HCs
RGCs
UMAP_1
UMAP_2
putative Ciliary Margin (pCM)
Retinal Progenitor Cells (RPCs)
Neurogenic RPCs (NRPCs)
Muller glia cells (MCs)
T1
T2
T3
Photoreceptor precursors (PPs)
Cone precursors (Cone-Ps)
Cones
Rods
Bipolar cells (BCs)
HC/AC/RGC progenitors (HC/AC/RGC-P)
Horizontal cells (HCs)
Amacrine cells (ACs)
Retinal ganglion cells (RGCs)
Non-Retinal
RPE
Ocular vesicle epithelium (OV-Epi)
Ocular surface epithelium progenitors (OSE-P)
Ocular surface epithelium (OSE)
Neuronal progentitors (NPs)
Neural crest derived cells (NCs)
Corneal stroma (CSt)
Corneal endothelium (CE)
Corneal fibroblasts (CF)
Fibroblasts (F)
Astrocytes (A)
Pluripotent stem cells (PSCs)

**Figure S7: ScATAC-Seq of ROs during early (day 10 - 35) and mid (day 45 - 90) developmental stages**. **A-C)** UMAP plots of scATAC-Seq of ROs at day 10 (**A**), day 20 (**B**) and day 35 (**C**) of differentiation. **D-E)** scATAC-Seq UMAP plots of ROs at day 45 (**D**), day 60 (**E**) and day 90 (**F**). Cluster identity defined on the basis on gene activity scores, calculated based on open regions of the chromatin of retinal specific cell markers (Table S5).

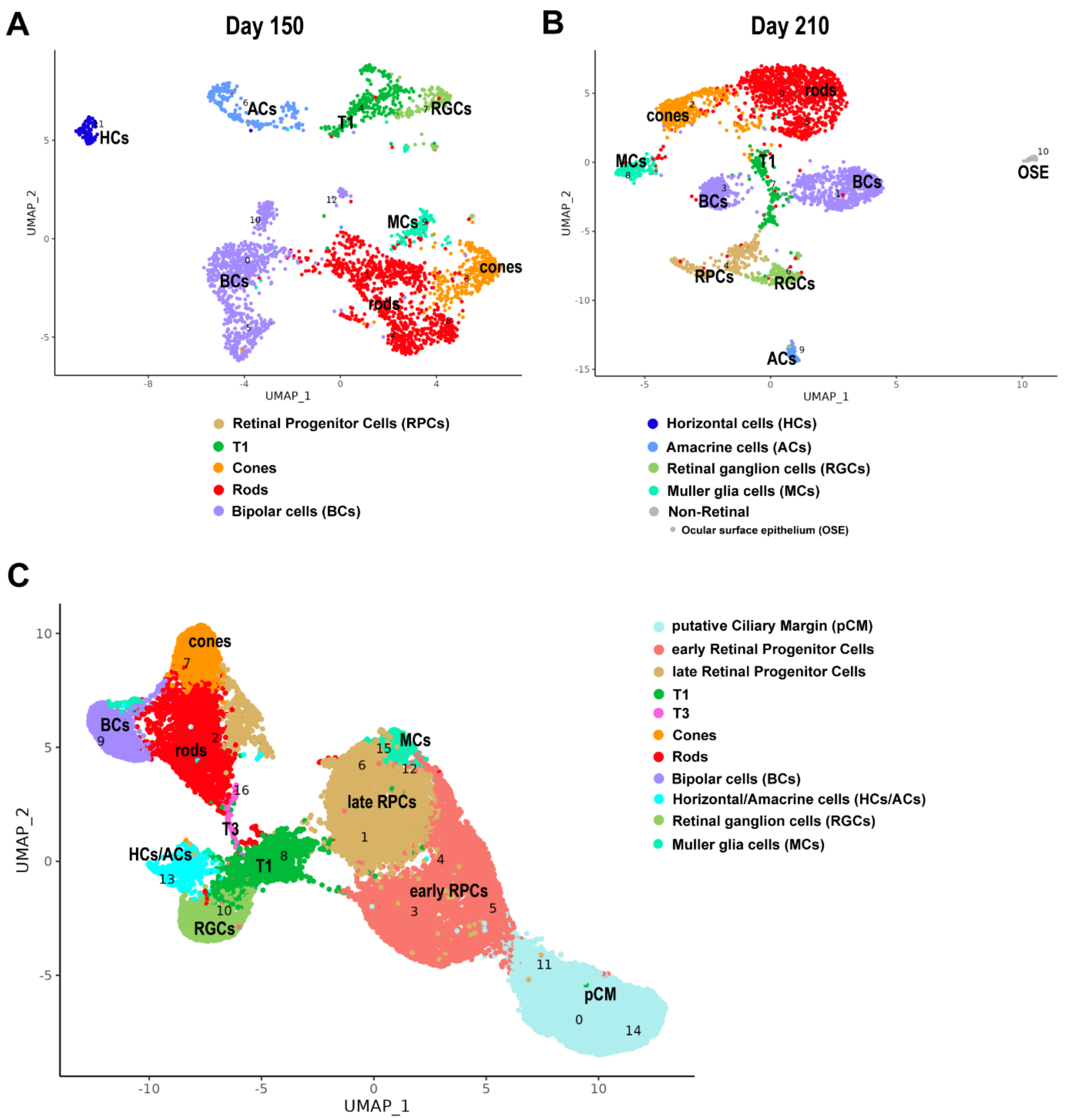


**Figure S8: ScATAC-Seq of ROs during late development (day 150 - 210) and integrated ROs UMAP. A, B)** UMAP plots of scATAC-Seq of ROS at day 150 (**A**) and day 210 (**B**). **C)** Integrated ATAC-Seq data of ROs from day 10 to day 210 of differentiation. Cluster identity defined based on gene activity scores, calculated based on open regions of the chromatin, of retinal specific cell markers (Table S5).

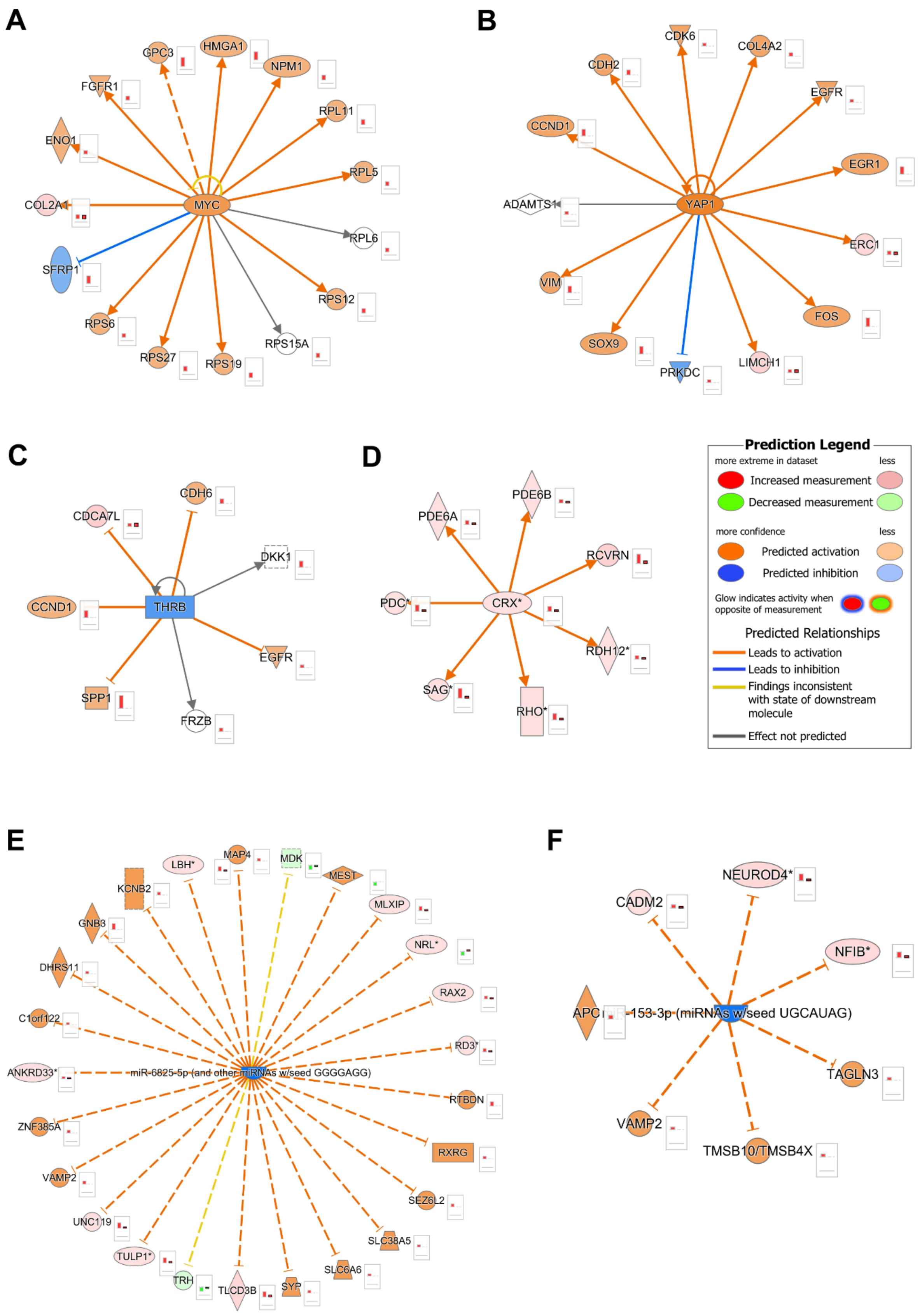


**Figure S9: Representative gene regulatory networks in early (A) and late RPCs (B, C), photoreceptors (D, E), and BCs (F) depicting activated and inhibited upstream regulators and their target genes.** IPA was used to generate the upstream regulatory networks from differentially expressed genes from the scRNA-Seq data and differential accessibility analysis in the scATAC-Seq data (Table S8). These networks indicate predictions of upstream regulators which might be activated or inhibited to explain observed upregulation/downregulations in the data. The bar plots next to each molecule illustrate the relative expression in the scRNA-Seq (column 1) and scATAC-Seq (column 2) datasets. The network nodes/bar plots colours represent observed upregulation/ increased chromatin accessibility (red), predicted upregulation/increased chromatin accessibility (orange), observed downregulation (green) and predicted downregulation/ decreased chromatin accessibility (blue). The edge colour represents the relationships between the molecules; orange = prediction and observation are consistent with activation; blue = prediction and observation are consistent with downregulation; yellow = prediction and observation are inconsistent; and grey relationship between the molecules is not available in the IPA knowledge database. *- indicates duplicates in scATAC-Seq data.

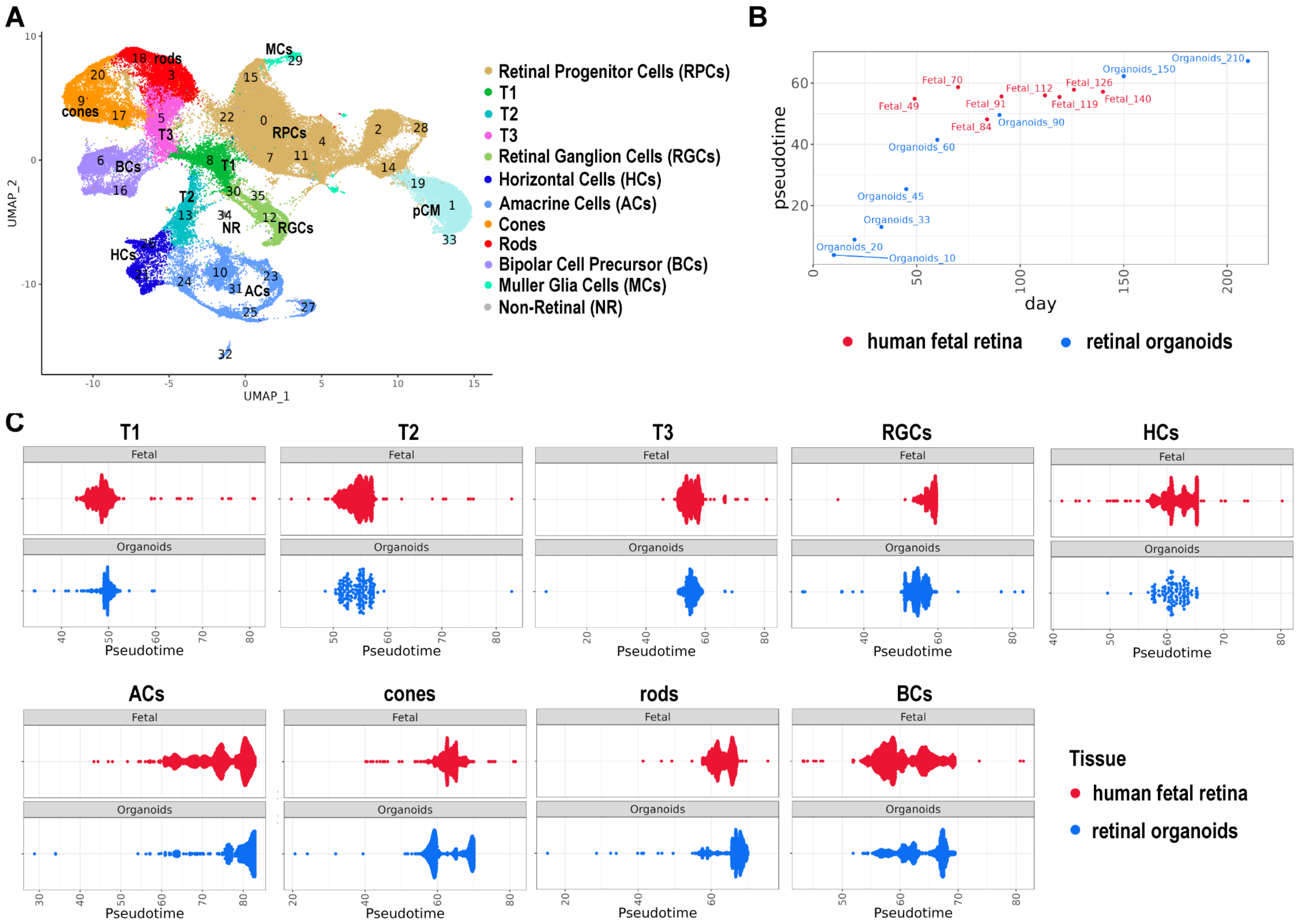
A
B
C
Retinal Progenitor Cells (RPCs)
T1
T2
T3
Retinal Ganglion Cells (RGCs)
Horizontal Cells (HCs)
Amacrine Cells (ACs)
Cones
Rods
Bipolar Cell Precursor (BCs)
Muller Glia Cells (MCs)
Non-Retinal (NR)
UMAP_1
UMAP_2
pseudotime
day
human fetal retina
retinal organoids
T1
T2
T3
RGCs
HCs
ACs
cones
rods
BCs
Fetal
Organoids
Pseudotime
Tissue
human fetal retina
retinal organoids

**Figure S10: Pseudotime trajectory comparison of chromatin accessibility landscapes of ROs and developing human retina indicates minor differences in the temporal emergence and abundance of the retinal neurons. A)** Integrated UMAP of ROs (day 10 – 210) and human fetal retina (8 – 21 post conception weeks) scATAC-Sec datasets (Table S9). **B)** Grouping of individual developmental stages of ROs (blue) and human fetal retina samples (red) within pseudotime analysis, indicating that RO differentiation ROs closely follows the retinal development *in vivo*. **C)** Pseudotime trajectory analysis of individual cell types demonstrate some differences in T1, RGCs, cone, rod, and BCs emergence between ROs and human fetal retina.

**Table S1 (separate file)**
**ScRNA-Seq analysis of ROs at day 10 – 210 of differentiation.** Cluster annotations are based on highly expressed markers for each cell type and are shown on separate spreadsheets for each sample.

**Table S2 (separate file)**
**Spatial transcriptomic analysis of ROs at day 10 – 210.** Cluster annotations are based on highly expressed markers for each cell type and are shown on separate spreadsheets for each sample.

**Table S3 (separate file)**
**Single cell RNA-Seq analysis of eye-like organoids at day 45 and day 90 of differentiation.** Cluster annotations are based on highly expressed markers for each cell type and are shown on separate spreadsheets for each sample.

**Table S4 (separate file)**
**Integrated scRNA-Seq analysis of ROs (day 10 – 210 of differentiation) and pseudotime analysis.** The first spreadsheet shows the integrated scRNA-Seq data for ROs (day 10 – 210 of differentiation). The second spreadsheet displays highly expressed markers characterising lineage transitions from pCM, RPCs to T1, T2, T3. The third spreadsheet shows highly expressed markers characterising lineage transitions from pCM, RPCs, T1 to T3, photoreceptors and bipolar cells. The fourth spreadsheet shows highly expressed markers characterising lineage transitions from pCM, RPCs to T1, T2, RGCs, horizontal and amacrine cells.

**Table S5 (separate file)**
**ScATAC-Seq analysis of ROs at day 10 – 210 of differentiation**. Gene activity estimates were generated using Signac. Genes with high activity scores for each cell type are shown on separate spreadsheets for each sample. The final spreadsheet shows the gene activity scores for the integrated datasets. All retinal cells across samples were integrated using harmony and the data was then re-clustered to produce integrated results.

**Table S6 (separate file)**
**Differential accessibility peaks for each retinal cell type using the integrated scATAC-Seq dataset.** Peaks were linked to genes using Cellranger and classified as either promoter, distal or intergenic. The distance from the gene is shown for each peak. Additional abbreviations: eRPCs – early RPCs, lRPCs – late RPCs, HCs- horizontal cells, ACs- amacrine cells.

**Table S7 (separate file)**
**Predicted transcription factor binding (motifs) for each cell type in ROs using the integrated scATAC-Seq dataset**. ChromVAR was used to characterise transcription factor motifs for each cell type. Additional abbreviations: eRPCs – early RPCs, lRPCs – late RPCs, HCs- horizontal cells, ACs- amacrine cells.

**Table S8 (separate file)**
**List of significant gene expression regulators in RPCs, transient neurogenic progenitors (T1, T2), photoreceptors, bipolar, horizontal and amacrine cells.**

**Table S9 (separate file)**

**Integrated data set from scATAC-Seq of ROs (day 10- 210 of differentiation) and human developing retina (pcw 10 – 21).** Peaks were linked to genes using Cellranger and further processed with Signac. Harmony was used to reduce batch differences between ROs and human developing retina samples.
